# Phylogenomics of scorpions reveal a co-diversification of scorpion mammalian predators and mammal-specific sodium channel toxins

**DOI:** 10.1101/2020.11.06.372045

**Authors:** Carlos E. Santibáñez-López, Shlomi Aharon, Jesús A. Ballesteros, Guilherme Gainett, Caitlin M. Baker, Edmundo González-Santillán, Mark S. Harvey, Mohamed K. Hassan, Ali Hussin Abu-Almaaty, Shorouk Mohamed Aldeyarbi, Lionel Monod, Andrés Ojanguren-Affilastro, Robert J. Raven, Ricardo Pinto-Da-Rocha, Yoram Zvik, Efrat Gavish-Regev, Prashant P. Sharma

**Affiliations:** Department of Integrative Biology, University of Wisconsin–Madison, Madison, WI 53706; National Natural History Collections, The Hebrew University of Jerusalem, Jerusalem, Israel; Department of Organismic and Evolutionary Biology, Harvard University, Cambridge, MA 02138; Laboratorio de Aracnología, Facultad de Ciencias, Departamento de Biología Comparada, Universidad Nacional Autónoma de México, C.P. 04510 Mexico City, Mexico; Department of Terrestrial Zoology, Western Australian Museum, Locked Bag 49, Welshpool DC 6986, Western Australia, Australia; Zoology Department, Faculty of Science, Port Said University, Egypt; Département des arthropodes et d’entomologie I, Muséum d’histoire naturelle, Route de Malagnou 1, 1208 Geneve, Switzerland; División Aracnología, Museo Argentino de Ciencias Naturales, Av. ángel Gallardo 470, Buenos Aires, Argentina; Department of Terrestrial Biodiversity, Queensland Museum, Grey Street, South Bank, Brisbane, Queensland, Australia; Departamento de Zoologia, Instituto de Biociências, Universidade de São Paulo, Rua do Matão, travessa 14, 321, São Paulo, Cep: 05508-900, Brasil; Hoopoe Ornithology & Ecology Center, Yeroham, Israel & Ben-Gurion University of the Negev, Be’er-Sheva, Israel

**Keywords:** Arachnida, venom, phylostratigraphy, dating, co-diversification

## Abstract

Scorpions constitute a charismatic lineage of arthropods and comprise more than 2,500 described species. Found throughout various tropical and temperate habitats, these predatory arachnids have a long evolutionary history, with a fossil record that began in the Silurian. While all scorpions are venomous, the asymmetrically diverse family Buthidae harbors nearly half the diversity of extant scorpions, and all but one of the 58 species that are medically significant to humans. Many aspects of scorpion evolutionary history are unclear, such as the relationships of the most toxic genera and their constituent venom peptides. Furthermore, the diversification age of toxins that act specifically on mammalian ion channels have never been inferred. To redress these gaps, we assembled a large-scale phylogenomic dataset of 100 scorpion venom transcriptomes and/or genomes, emphasizing the sampling of highly toxic buthid genera. To infer divergence times of venom gene families, we applied a phylogenomic node dating approach for the species tree in tandem with phylostratigraphic bracketing to estimate minimum ages of mammal-specific toxins. Our analyses establish a robustly supported phylogeny of scorpions, particularly with regard to relationships between medically significant taxa. Analysis of venom gene families shows that mammal-specific sodium channel toxins have independently evolved in five lineages within Buthidae. The temporal windows of mammal-specific toxin origins are contiguous with the basal diversification of major scorpion mammal predators such as carnivores, shrews, bats and rodents. These results suggest an evolutionary arms race model comprised of co-diversification of mammalian predators and NaTx homologs in buthid venom.

---

To scientists and laypersons, scorpions are fascinating for the potency of their venom, a complex mixture of bioactive compounds secreted in specialized organs in the ultimate body segment, and used to disrupt biochemical and physiological processes in target organisms (King and Hardy 2013; Casewell et al. 2013). This lineage of arachnids diversified in the Permian and harbors considerable modern diversity; nearly 2500 species have been described in the presently 22 recognized families, with this diversity distributed across all continents except Antarctica (Sissom 1990; Santibáñez-López et al. 2019a, b; Tropea and Onmis 2020). Scorpions are particularly well-adapted to survival in extreme habitats, and their ability to produce and deliver venoms is inferred to be a key factor to their success. A major dimension of their biodiversity is found in their venoms, which are rich in toxins that have a broad array of biological targets (including those affecting Na^+^, K^+^, Cl^−^ and Ca^2+^ ion channels), mucopolysaccharides, and enzymes (Cao et al. 2013; He et al. 2013). Scorpion envenomation causes nearly an order of magnitude greater fatalities worldwide than snakebites, and particularly so in developing or rural subtropical regions. Intensive functional study of specific peptides has uncovered significant biomedical applications in scorpion venom, such as the identification of antimicrobial and antitumor agents, fluorescent “tumor paint”, and transport molecules for molecular cargo (Veiseh et al. 2007; Rapôso 2017; Díaz-Perlas et al. 2018).

While all scorpions are venomous, both the extant diversity of scorpions, as well as the toxicity of their venom to mammals, is asymmetrical distributed; ca. 1200 species (47% of described diversity) of scorpions are members of the family Buthidae (“thick-tailed scorpions”; Fig. 1), which includes nearly all significantly venomous scorpion species, as well most of the known molecular diversity of scorpion venom (Santibáñez-López et al. 2019b). All scorpions are thought to possess insect-specific ion channel toxins, which facilitate prey capture. Within buthids, salient components of the venom cocktail are ion channel toxins that are specific to mammalian targets and are inferred to function as anti-predator deterrents (Niermann et al. 2020). Such neurotoxins operate by blocking action potentials at nerve synapses, precipitating symptoms of neurotoxicosis such as intense pain, hypersalivation, muscle spasms, asphyxia, and paralysis. The genera Androctonus, Buthus, Centruroides, Hottentotta, Leiurus, Parabuthus and Tityus all contain multiple highly toxic and medically significant species known for the potency of their neurotoxins (Santos et al. 2016; Niermann et al. 2020).

**Figure 1.**
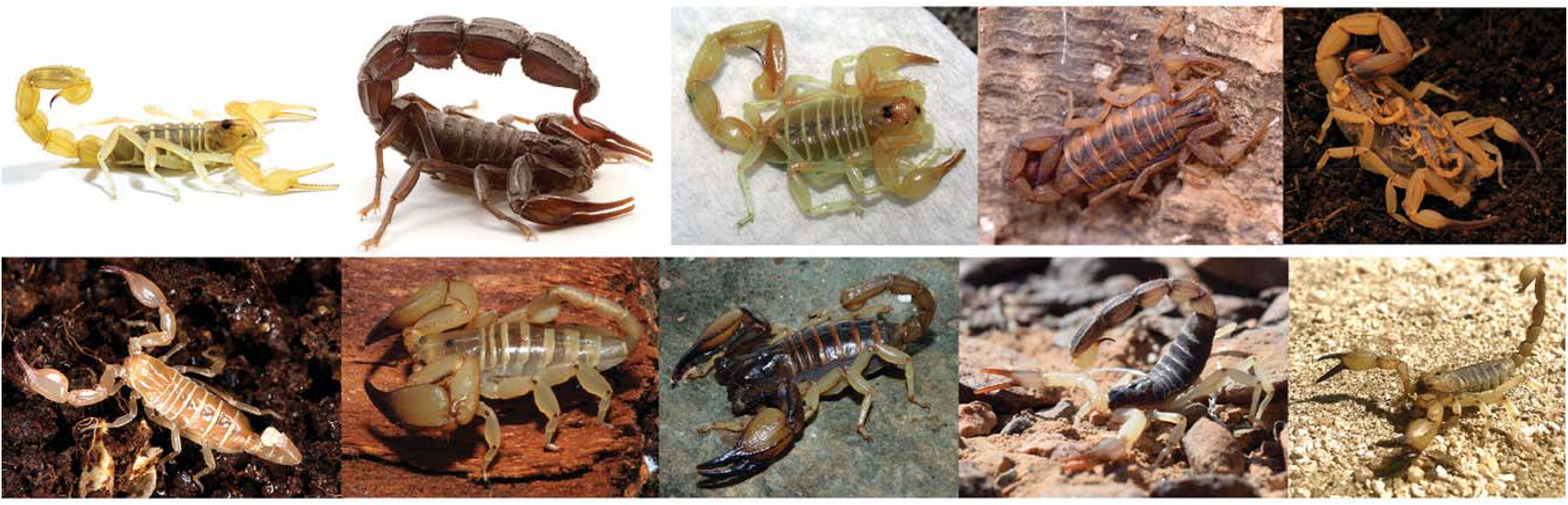
Exemplars of scorpion diversity in the two major parvorders. Top row, buthid species. Bottom row, Iurida species. Photos: *Buthus israelensis* and *Androctonus crassicauda* (R. Livne); *Buthacus leptochelys*, *Palaeocheloctonus pauliani* and *Opistophthalmus carinatus* (J. Ove Rein); *Centruroides meisei* and *Tityus serrulatus* (B. Myers); *Belisarius xambeui* (G. Giribet); *Hadrurus obscurus* and *Anuroctonus bajae* (C. Santibáñez-López). All photographs published with permission.

Surprisingly, little is understood as to about the evolutionary relationships of medically relevant scorpions. The higher-level molecular phylogeny of buthids was first inferred using a 296-bp fragment of 16S rRNA and sampling 17 genera, with weak resolution of basal relationships (Fet et al. 2003). Subsequently, most molecular phylogenetic studies of Buthidae have used a handful of Sanger-sequenced loci to address relationships of derived groups, such as individual genera or subfamilies, or of buthids restricted to specific geographic terranes (Ojanguren-Affilastro et al. 2017; Suranse et al. 2017). Mitochondrial DNA surveys of medically significant buthids in particular have not yielded supported basal relationships (Borges et al. 2014). Furthermore, in the first phylogenomic study of scorpions, sampling of Buthidae was limited to just four species (Sharma et al. 2015). While subsequent analyses of venom evolution added buthid terminals to the scorpion tree, these additional datasets were derived from Sanger-sequenced EST libraries or 454-pyrosequenced datasets, resulting in high proportions of missing data and attendant instability in buthid relationships (Santibáñez-López et al. 2018). Other recent phylogenomic investigations have prioritized the systematically complex Iurida, without adding to the sampling of the original four Buthidae transcriptomes (Sharma et al. 2018; Santibáñez-López et al. 2019a, b; Santibáñez-López et al. 2020).

The lack of a robust and densely sampled molecular phylogeny of scorpions has hindered various evolutionary analyses pertaining to the evolution of scorpion morphology, biogeography, and venomics. In particular, undersampling of buthids risks omitting key nodes for understanding the mode and tempo of scorpion and scorpion venom evolution. Targeting this group in phylogenomic analyses is key to testing hypotheses about single versus multiple origins of mammal-specific toxins, as well as their potential co-diversification with mammalian predators. To close these knowledge gaps, we assembled a phylogenomic dataset of 100 scorpion terminals. Within Buthidae, we emphasized sampling of highly toxic species from fieldwork theaters in the southwestern US, the Neotropics, and the Middle East. Here, we show that mammal-specific toxins have originated independently in five major clades within Buthidae and the timing of their origins coincides with the diversification of major mammalian predators of scorpions.

## Materials and Methods

Extended methods are provided in the Supplementary Text.

### Fieldwork, RNA-Seq, and Phylogenomic Analyses

Scorpions were collected by hand in field theaters across Brazil, Egypt, Israel, and the US, commonly with the aid of ultraviolet lighting. A subset of well-studied species was obtained through captive breeding programs. Milking and dissection of venom glands, RNA extraction, and paired-end transcriptome sequencing was performed on the Illumina HiSeq 2500 platform for 42 species, following our previous approaches (Sharma et al. 2015; Santibáñez-López et al. 2018). New datasets were combined with 45 venom gland RNA-Seq datasets and one genome we previously generated (Santibáñez-López et al. 2018; Schwager et al. 2017). Twenty outgroup species were including in the analysis, spanning tetrapulmonates, pseudoscorpions, harvestmen, and three horseshoe crab genomes. All collecting, vouchering, and accession data are provided in Supplementary Tables S1-4.

Orthologous loci were drawn from MCL clustering of 3564 orthogroups computed from a larger analysis of Chelicerata and outgroup taxa (Ballesteros and Sharma 2019). Untrimmed alignments were used to produce a hidden Markov profile using hmmerbuild from hmmer package v.3.2.1 (Mistry et al. 2013). Each proteome/transcriptome of the species of interest was then searched (hmmerseacrh) for matches against the collection of profiles with an expectation threshold of e < 10^−20^; for cases with more than one hits per locus, the sequence with the best score was selected, and the corresponding sequence appended to the locus FASTA file aggregating the putative orthologous found in each species. Clustering of putative orthologs was tested by comparing each orthogroup’s constituent sequences with the proteome of the fruit fly *Drosophila melanogaster*, and removing from orthogroups any sequences with mismatching functional annotations. Finally, alignments and gene trees were visually inspected for evident paralogy. Paralogous sequences were discarded; if the number of sequences retained post-trimmed was below the minimum taxon occupancy threshold, the entire locus was discarded. Three matrices were assembled with minimal taxon occupancy thresholds: Matrix 1 (at least 115 species), Matrix 2 (at least 109 species), and Matrix 3 (at least 103 species). Phylogenetic inference of these concatenated matrices was computed with IQ-TREE v. 1.6 (Nguyen et al. 2014) implementing the best-fitting amino acid substitution model per partition (-spp; Supplementary Text). As approaches based on concatenation can be prone to incorrect inference, species trees were estimated using the coalescent method implemented in ASTRAL v.3 (Mirarab and Warnow 2015), using the collection of orthologous gene trees as inputs. Analysis of the smallest matrix (Matrix 3) was trialed using Phylobayes-mpi v. 1.7 (Lartillot and Philippe 2004) with four independent chains under the CAT + GTR + Γ_4_ model.

### Divergence Time Estimation

Divergence time estimation was computed on Matrix 2 using the approximate likelihood calculation as implemented in Codeml and MCMCtree (both part of the PAML v. 4.8 software package (Yang 2007; dos Reis and Yang 2019). The ML tree inferred from Matrix 2 was used as the input tree and was calibrated using fifteen fossil taxa (Supplementary Text, Supplementary Table S5). Four Bayesian inference chains were run for 2.5 M post-burnin generations (burnin of 25000 generations); convergence diagnostics were assessed using inbuilt tools in MCMCTree (Puttick 2019).

### Toxin Homology and Evolution

Cysteine-stabilized α-helix and β-sheet fold (CSαβ), disulphide-directed beta-hairpin (DDH) and Inhibitor cystine knot (ICK) homologs from scorpion venom were retrieved from the complete dataset used in the scorpion phylogenetic analyses following our recent approaches (Santibáñez-López et al. 2018; Santibáñez-López et al. 2019b), and from UniProt (Supplementary Table S6). Gene trees were conducted using IQ-TREE for the entire dataset (1,353 CSαβ-ICK scorpion toxins, with 41 DDH scorpion toxins as outgroups), and for each of the four main clades recovered: (a) sodium channel toxins (NaTx); (b) potassium channel toxins (KTx); (c) chlorine channel toxins (ClTx); and (d) calcins. Comparative analyses between the subclades recovered within the NaTx included the search for repetitive motifs in their mature peptide using Multiple Em for Motif Elicitation (Bailey et al. 2015). The mature peptide of the two main clades within the NaTx (Aah2-like and Cn2-like) were separately analyzed using CLANS clustering (Frickey and Lupas 2004). Estimation of minimum ages for mammal-specific toxins was derived from molecular dating performed herein, using the ages of the most inclusive clades of taxa as minimum age estimates for gene age (phylostratigraphic bracketing).

Divergence times for scorpion predators such Herpestidae (Carnivora), Chiroptera, Eulipotyphla and Rodentia were retrieved from a recent analysis of mammal diversification times (Upham et al. 2019) and compared against the mammal-specific toxin origins.

## Results

### Phylogenomic analysis

To infer the phylogenomic relationships of scorpions, we generated a dataset comprised of 99 venom gland transcriptomes (97 sequenced by us) and the genome of the buthid *Centruroides sculpturatus* (Schwager et al. 2017), with 42 new assemblies sequenced herein (Supplementary Tables S1-4). Inclusion of an older pyrosequenced genome of *Mesobuthus martensii* and Sanger-sequenced EST libraries from other buthid species was trialed as separately from main analyses to the degree of missing data incurred by these terminals. Outgroup taxa consisted of 20 transcriptomes spanning seven non-scorpion chelicerate orders. Inference of orthology followed a previously established pipeline, drawing upon groupings of MCL clusters established in a previous analysis of chelicerate relationships, followed by phylogenetically informed identification of orthologous groups (Ballesteros and Hormiga 2016; Ballesteros and Sharma 2019). Three phylogenomic matrices were constructed spanning 192 to 660 loci (53,333 to 185631 aligned amino acid sites), using taxon minimum occupancy thresholds of 103, 109, and 115 terminals per locus (88.0%, 91.4%, and 94.1% complete, respectively). We refer to these henceforth as Matrices 1-3, respectively.

Maximum likelihood analyses of concatenated datasets using IQ-TREE v. 1.6 consistently recovered with maximal nodal support the monophyly of scorpions and the established basal split between Buthida (consisting of Buthidae, Chaerilidae, and Pseudochactidae) and Iurida (the remaining scorpion families) (Sharma et al. 2015). Matrices 1 and 2 recovered support for *Lychas variegatus* as the sister group of the remaining buthids (87-95% bootstrap frequency), whereas Matrix 3 recovered a nested placement of *L. variegatus* with weak nodal support (72%; Supplementary Figs. S1-S3). All three datasets recovered with maximal support four major clades within the buthids, consisting of (a) the largely Palearctic “*Buthus* group” (*Androctonus*, *Birulatus*, *Buthus, Buthacus*, *Compsobuthus*, *Hottentotta*, *Leiurus*, and *Orthochirus*, with *Mesobuthus* recovered in this group in supplementary analyses), (b) *Ananteris* and *Babycurus*, (c) the “*Uroplectes* group” (*Grosphus*, *Parabuthus* and *Uroplectes*), and (d) the “*Tityus* group” (*Centruroides*, *Heteroctenus*, *Jaguajir, Physoctonus, Rhopalurus*, *Tityus,* and *Troglorhopalurus*). Relationships between these groups across all supermatrix analyses supported the sister group relationship of the *Tityus* and *Uroplectes* groups, with this lineage in turn sister group to (*Ananteris* + *Babycurus*). Relationships within Iurida closely reflected tree topologies reported in our previous works and are not discussed further herein (Sharma et al. 2015; Santibáñez-López et al. 2019a, b; Santibáñez-López et al. 2020).

In addition to maximum likelihood tree reconstruction based on a supermatrix approach, we inferred relationships using multispecies coalescent (MSC) tree reconstruction using ASTRAL v.2 for the constituent gene trees of Matrices 1-3. Higher-level relationships of scorpions were largely congruent across analyses, excepting the placement of *Lychas*, which was unsupported (posterior probability [PP] = 0.47-0.80 across analyses; Supplementary Fig. S4).

While we trialed the use of Phylobayes-mpi to estimate phylogenomic relationships under the CAT + GTR + Γ_4_ model, Bayesian inference chains consistently failed to converge (*maxdiff* = 1.0) after 10,596 −11,574 cycles of computational effort across four chains, equaling six months of continuous wall-clock computation. The resulting tree topology of the chain with the highest log-likelihood nevertheless reflected the same relationships as the maximum likelihood inferences (Supplementary Fig. S5).

### Molecular dating

Phylogenomic estimation of divergence times was inferred using a node dating approach with MCMCTree on Matrix 2, implementing a likelihood approximation of branch lengths using a multivariate normal distribution (Yang 2007; dos Reis and Yang 2019). Fossils used to inform the dating consisted of six ingroup and nine outgroup node calibrations. All calibrations were implemented as soft minimum and soft maximum ages. Identity of fossil taxa and implementation of temporal ranges as calibrations are provided in the Supplementary Text, and Supplementary Table S5. Divergence times were inferred under two models of rate evolution, a correlated rates model and an independent rates model. Both clock models recovered comparable inferences of basal scorpion diversification, with a split between Buthida and Iurida dating to the Carboniferous-Permian (291-320 Mya; highest posterior density [HPD] interval: 247-352 Mya) (Fig. 2). The diversification of Buthidae was estimated to span the end-Jurassic to the Early Cretaceous (95% HPD: 105-161 Mya). Divergences within the major buthid clades fell within the Late Cretaceous to Paleogene, excepting the *Buthus* group, wherein most divergences were estimated to occur in the Neogene.

**Figure 2.**
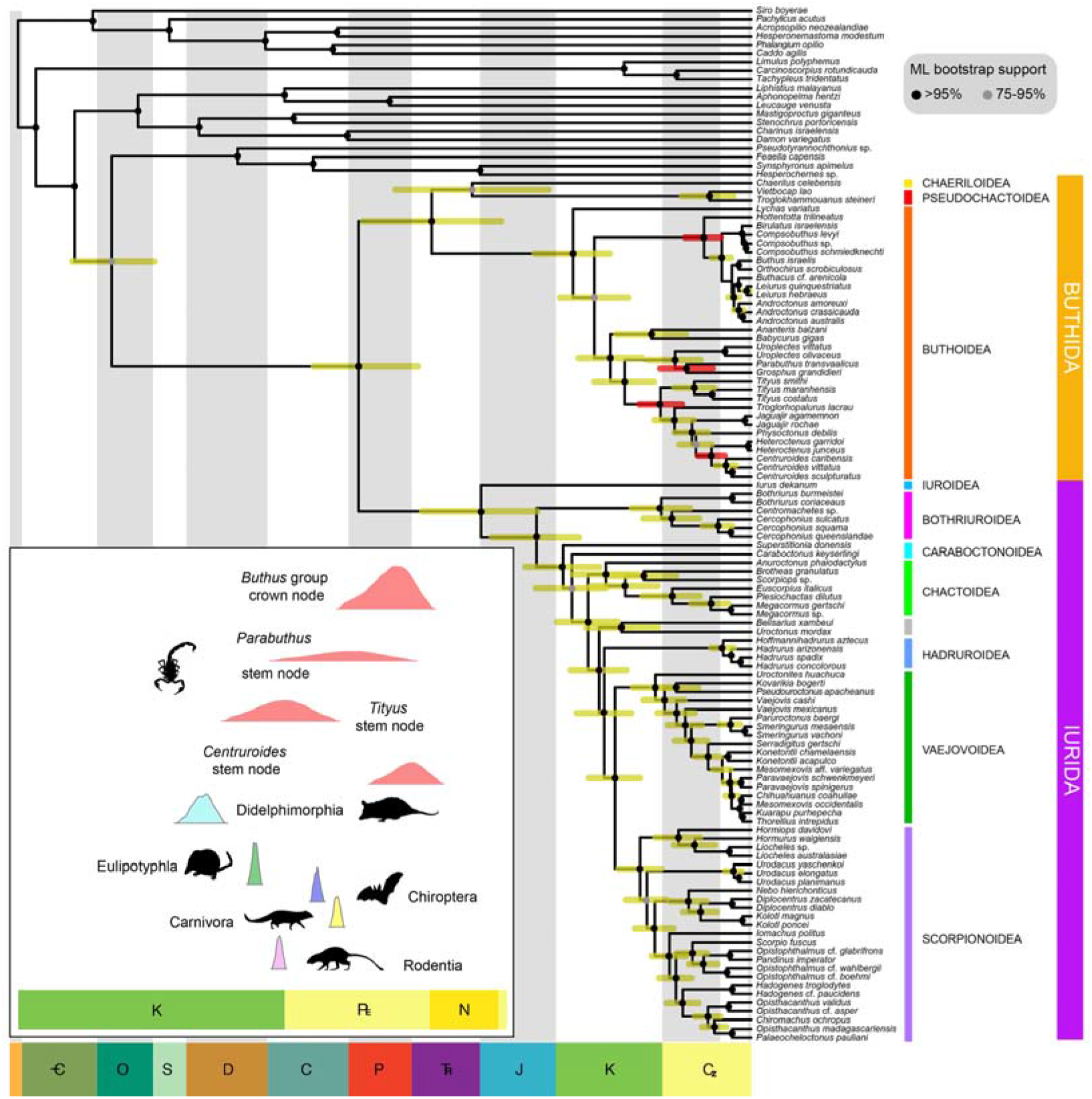
Chronogram of scorpion relationships derived from maximum-likelihood analysis of 192 loci (53,333 amino acid sites). Node ages are computed in a time-calibrated analysis using 15 fossil calibrations. Blue bars depict 95% credibility intervals of node ages, whereas red bars depict 95% credibility intervals for the most toxic buthids. Inset: Posterior distribution of node ages corresponding to mammal-specific toxin origins (in red), compared to the posterior distribution of the four major mammal orders that include scorpion predators. Mammal order ages from (43).

### Phylostratigraphic bracketing of venom gene family origins

Putative cysteine-stabilized α-helix and β-sheet fold (CSαβ) and Inhibitor cystine knot (ICK) venom components were identified in the 100 terminals and from the UniProt databases. Within these peptides, the most relevant are those affecting the ion channels, such as the sodium channel toxins (NaTx), potassium channel toxins (KTx), chlorotoxins (ClTx), and the ryanodine receptor ligands (Calcins). Using gene tree reconstructions, we documented 1,353 CSαβ-ICK peptide homologs spanning 151 scorpion species in 19 families (from which 48% were buthids; Supplementary Table S6). Maximum likelihood analysis of the CSαβ-ICK matrix and 41 disulphide-directed beta-hairpin (DDH) sequences as outgroups (399 amino acid sites) recovered a gene tree subdivided into three major clades: (a) NaTx; (b) the calcins; and (c) the KTx including the nested ClTx (Supplementary Fig. S6-7). Our results reveal that calcins are phylogenetically restricted to iurids (Supplementary Figs. S6, S8), whereas ClTx are restricted to a subset of Old World buthids (Supplementary Figs. S6, S9). Within KTx, ML analysis recovered the presence of nine clusters, corresponding to (a) αKTx; (b) βKTx; (c) εKTx (unique to buthids); (d) κKTx (unique to iurids); (e) λKTx (unique to buthids); (f, g) Scorpine-like type 1 and type 2; and (h, i) Kunitz-type 1 and type 2 (Supplementary Fig. S7).

Within NaTx, mammal-specific toxins were recovered as six clusters within two major clades: (a) the Aah2-like; and (b) the Cn2-like clades (Figs. 3A-B). The Aah2-like gene family was found exclusively in Buthida (mostly from Old World Buthids, Supplementary Fig. S10-11) and encodes for peptides with arthropod affinity, insect and mammal affinity, and mammal-affinity (Figs. 3B). In contrast, within the Cn2-like gene family, we found one cluster restricted to Iurida, and six clusters exclusively in Buthida (Supplementary Figs. S11-15). Among these, two clusters with mammal-specific targets were found uniquely in the genera *Centruroides* and *Tityus* (Supplementary Figs. S14-15). Our search for motifs with MEME v. 5.1 (Bailey et al. 2015) showed a short non-unique motif of 10 amino acids (GXXCWCXXLPD) for members of the Aah2-like gene family, and no specific motif for members of the Cn2-like gene family. More specifically, no conserved or repetitive motifs were found in the mammal-specific toxins of either the Aah2 or Cn2 gene families (Supplementary Fig. S16). Given the short sequence length of NaTx, we also assessed patterns of relatedness between the 661 mature peptide sequence from our dataset using CLANS clustering (Frickey and Lupas 2004). Consistent with the gene tree analyses, this approach recovered seven major groups of peptides with mammal-specific affinity (Fig. 3C).

**Figure 3.**
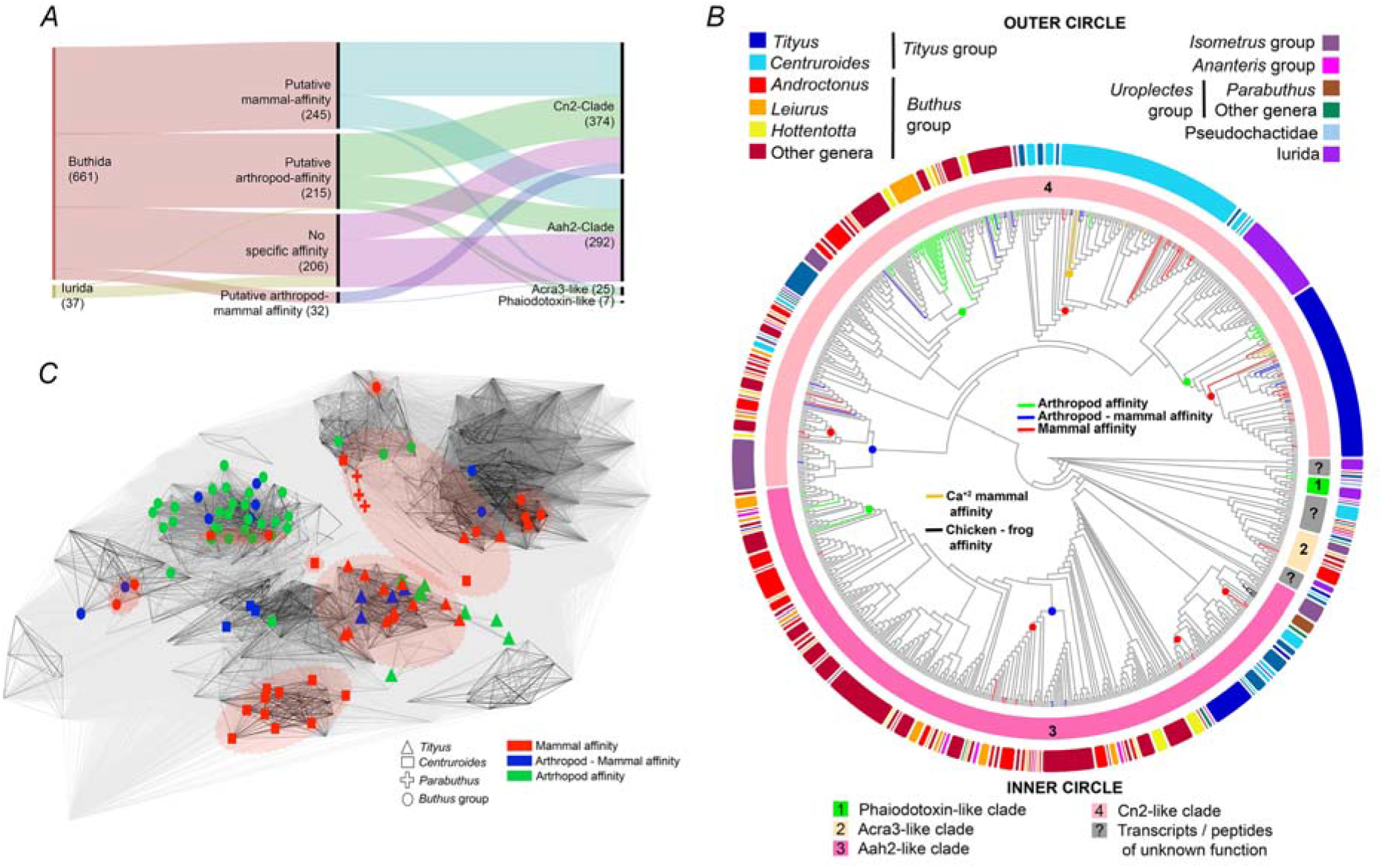
Evolutionary analyses of the sodium channel toxin family (NaTx). *(A)* Alluvial plot summarizing the affinity of the NaTx found in scorpion venom (center) of each parvorder (left), and their subtype (right). Numbers represent the total transcripts and/or peptides found in our transcriptomic analyses and UniProt. *(B)*. NaTx affinity (inner bars) plotted onto the NaTx gene tree subdivided into the seven subclades (four well known plotted onto the alluvial plot in panel A). In gray: transcript clades with unknown function. Circles in nodes represent the putative ancestral function as a result of the most parsimonious ancestral state reconstruction. (C) Three-dimensional CLANS clustering of the mature peptide amino acid sequence. Colors for NaTx affinity and lines indicating pairwise similarity. Red dotted lines show seven distinctive clusters that include at least one peptide with known mammal affinity.

To infer the age of mammal-specific genes, we employed a phylostratigraphic bracketing approach, using the stem age of the most inclusive buthid clade containing a given mammal-specific toxin as a proxy for the minimum estimated gene age. Stem ages of *Centruroides*, *Tityus*, *Parabuthus*, and the node uniting *Hottentotta* with the remaining *Buthus* group scorpions thus implied the earliest diversification of mammal-specific toxins in a temporal window spanning 20-83 Mya (Fig. 2, inset).

## Discussion

### Phylogenetic Relationships Within Buthidae Validate Morphology-Based Systematics

The systematic history of scorpions was previously largely dominated by morphological analyses, with marked contention between competing interpretations of homologies and cladistic practices (Sharma et al. 2015). Phylogenomic analyses initially refuted much of the traditional understanding of scorpion higher-level systematics, with analyses showing that morphological characters used to delimit families and superfamilial taxa are highly homoplastic and/or uninformative (Sharma et al. 2015). While scorpion classification was subsequently revised to bring taxonomic groupings into accordance with phylogenomic outcomes, a paucity of molecular datasets addressing higher-level buthid relationships has remained a hurdle for understanding the evolution of this asymmetrically large family.

Interestingly, this work support for generic groupings that were previously defined on the basis of positions of pedipalp trichobothria (sensory setae). Intensive surveys of trichobothrial position by Fet et al. (2005) were previously used to delimit six major groups of scorpions (the *Buthus*, *Ananteris*, *Tityus*, *Charmus*, *Isometrus*, and *Uroplectes* groups), with the Paleartic *Buthus* groups comprising the sister lineage of the remaining Buthidae. Our results accord precisely with the morphological conception of the *Buthus* group, the *Uroplectes* group, and the *Tityus* group, with some analyses additionally recovering the basally branching placement of the *Buthus* group, albeit with weak support (Supplementary Figs. S1, S4).

Placements of *Lychas*, *Ananteris*, and *Babycurus* do not accord exactly with the traditional morphological grouping (Supplementary Fig. S17). It was previously thought that *Lychas* fell within a poorly resolved “*Ananteris* group”, whereas *Babycurus* was thought to inhabit another group that includes *Isometrus* (the *Isometrus* group), a genus not sampled herein. However, the Paleotropical genera *Lychas* and *Isometrus* are likely closely related, as they are distinguishable mainly by the condition of the leg tibial spur (Koch, 1977). A previous phylogenetic study also recovered *Isometrus* as a close relative of *Lychas*, though with poor genomic representation of both taxa (Santibáñez-López et al. 2018).

Another group not sampled herein using transcriptomic data is the enigmatic *Charmus* group, which is geographically restricted to the Indian subcontinent, southeast Asia, and parts of East Africa. While molecular data are uncommon for this lineage, a separate analysis we performed— in which we combined Matrix 2 with available Sanger-sequenced data for some buthid lineages as well as a pyrosequenced genome for *Mesobuthus martensii*—also revealed that *Charmus* is part of the group that includes the Malagasy endemic genus *Grosphus* and the southern African genera *Parabuthus* and *Uroplectes* (Supplementary Fig. S17).

Our results thus differed with morphological delimitations of buthid relationships only with respect to the composition of the two groups that were also poorly resolved by the morphological data in that study (the *Isometrus* and *Ananteris* groups; Fet et al. 2005). These outcomes vindicate the phylogenetic utility of trichobothrial arrangement in establishing shallow-level relationships within scorpion families.

### Contemporaneous Diversification of Buthid Mammal-Specific Toxins and Scorpion Mammalian Predators

Previous genomic resources sampling scorpion venom diversity have focused on a handful of medically significant species, such as *Leiurus quinquestriatus* (Egyptian death stalker scorpion) and *Centruroides sculpturatus* (Arizona bark scorpion) (Veiseh et al. 2007; Borges and Graham 2014; Schwager et al. 2017). We focused herein on comparative analyses sampling venom gene expression broadly across scorpion phylogeny, toward characterizing the evolutionary dynamics that precipitated scorpion toxicity to mammals. We also endeavored to increase high-quality transcriptomic resources for species renowned for their toxicity in the genera *Androctonus*, *Centruroides*, and *Tityus*. Through these datasets, we discovered that certain classes of toxins are phylogenetically restricted; as examples, chlorotoxins occur only in Old World Buthidae; scorpines occur in all families except Buthidae; and calcins are restricted to Iurida (Supplementary Fig. S6).

Phylogenomic dating recovered a surprisingly young age for the basal diversification of Buthidae in the Late Mesozoic. We inferred 21 independent origins of sodium channel toxins with known specificity for mammalian targets spanning five separate buthid lineages, with three of these occurring in genera sampled with multiple terminals. Phylostratigraphic bracketing of these toxins’ origins recovered age estimates broadly overlapping the basal diversification dates of several mammal orders that include scorpion predators (Fig. 2). These mammal-specific buthid NaTx were nested within a cluster of NaTx that target arthropod tissues across all scorpions (Fig. 3).

These results are consistent with an evolutionary arms race, wherein mammal-specific toxins were derived in scorpions from insect-specific ancestral peptides, reflecting the derivation of an anti-predator defensive adaptation from peptides previously used to target prey (Fig. 4). Similar dynamics have recently been revealed in the highly venomous Australian funnel-web spiders, wherein δ-hexatoxins exhibit high evolutionary conservation, reflecting a defensive role for deterring vertebrate predators (Herzig et al. 2020). However, in scorpions, molecular signatures of selection revealed no consistent pattern of amino acid sequence evolution across groups of mammal-specific toxins, consistent with the inference of independent evolutionary origins of anti-vertebrate defensive peptides (Supplementary Fig. S14). As an extension of this arms race, counter-adaptations to scorpion venom are known to occur in some scorpion predators. For example, the grasshopper mouse *Onychomys torridus* exhibits reduced sensitivity to pain caused by the sting of the Arizona bark scorpion *Centruroides sculpturatus* (Rowe et al. 2013). The mechanism of this counter-adaptation was shown to be amino acid variants of a voltage-gated Na^+^ channel in *O. torridus* that have evolved to selectively bind *C. sculpturatus* toxins, thereby blocking action potential propagation. Parallel evolution of resistance to the venom of *C. sculpturatus* via modification of Na^+^ ion channels has also been suggested in the bat *Antrozous pallidus* (Hopp et al. 2017). Comparable molecular dynamics underlying the evolution of resistance to snake venom have evolved at least four times in mammal predators of snakes, such as mongooses and honey badgers (Barchan et al. 1992; Drabeck et al. 2015; Holding et al. 2016). Given the biodiversity of Buthidae and the molecular diversity of their venoms, broader, phylogenetically informed surveys of venom gland transcriptomics may uncover additional origins of mammal-specific toxin function in scorpions, as well as improve the precision of molecular divergence time estimation. Venom gland transcriptomic databases sampling poorly studied species may also offer additional targets for beneficial biomedical application that are not represented among established scorpion biomedical research programs.

**Figure 4.**
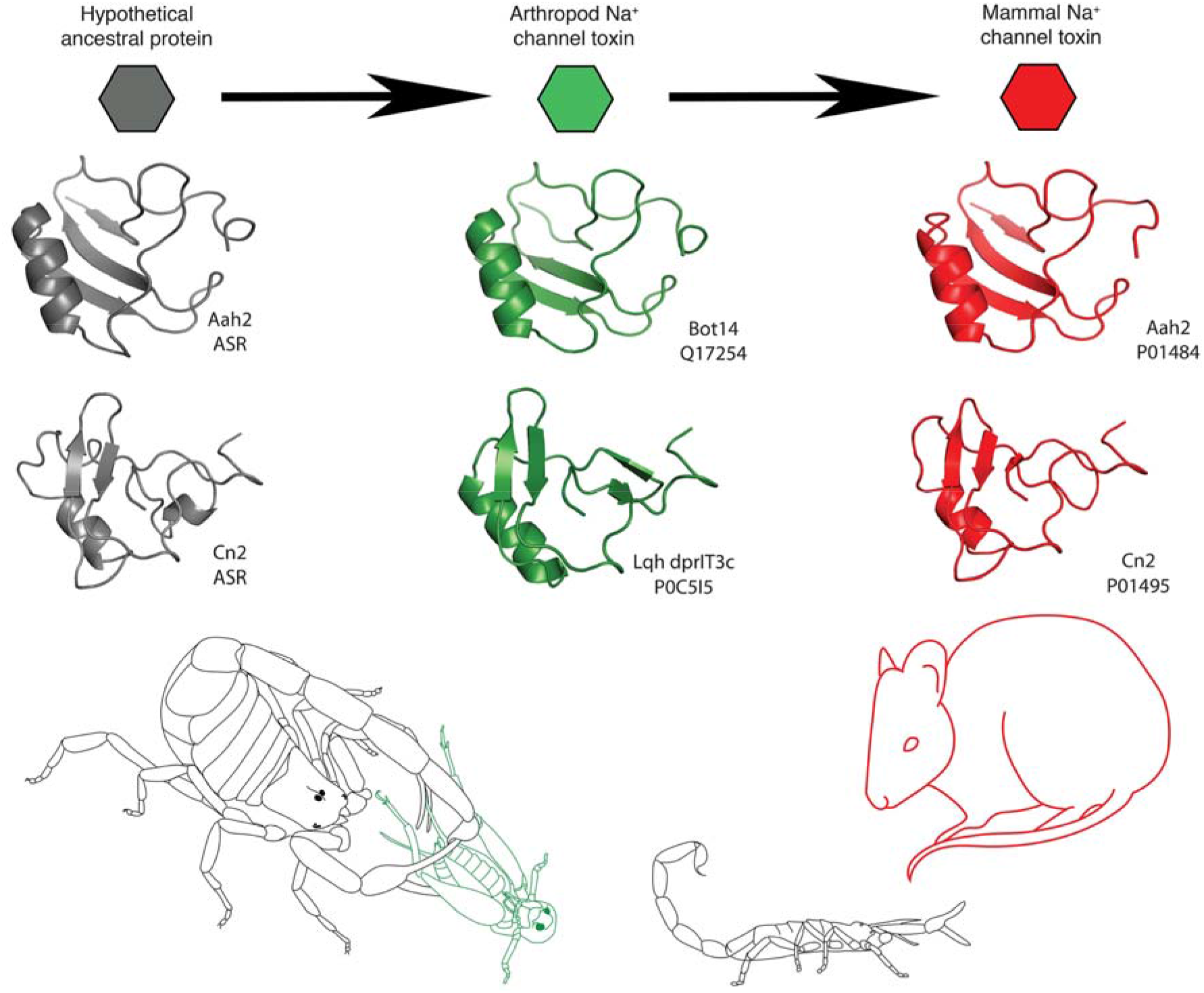
Secondary structures and proposed evolutionary pathway of scorpion Na+ channel toxins from the Aah (top model) and Cn2 (bottom model) clades. Our results suggest mammal specific toxins are derived from insect specific ones, which in turn evolved from ancestral molecules of unknown physiological activity. Mammal specific toxins from *A. australis* (pdb file 1ptx) and *C. noxius* (pdb file 1cn2) are shown in red; the model for the insect specific toxins from *B. tunetanus* (Q17254) and *L. hebraeus* (POC515) are shown in green; and, the hypothetical ancestral molecules of unknown function are shown in gray. The structures of these ancestral proteins are derived from the primary sequences estimated using Ancestral State Reconstruction.

## Acknowledgments

We are indebted to Brandon Myers for providing some of the specimens analyzed in this study. Steve Goodman kindly provided tissue of *Grosphus grandidieri* from Madagascar. Sequencing was performed at the UW-Madison Biotechnology Center. Access to computing nodes for intensive tasks was provided by the Center for High Throughput Computing (CHTC) and facilitated by Christina Koch; and by the Bioinformatics Resource Center (BRC) of the University of Wisconsin–Madison. Specimens in Israel were collected under permits 2017/41718 and 2018/42037, issued by the Israel National Parks Authority. Fieldwork in Israel was supported by a National Geographic Society Expeditions Council grant no. NGS-271R-18 to J.A.B. Fieldwork in Australia was supported by intramural UW-Madison funds. This work was supported by National Science Foundation (grant no. IOS-1552610) to P.P.S.; FAPESP (BIOTA, 2013/50297-0), National Science Foundation (DOB-1343578) to R.P.R.

## References

Ballesteros J.A., Hormiga G. 2016. A New Orthology Assessment Method for Phylogenomic Data: Unrooted Phylogenetic Orthology. Mol. Biol. Evol. 33:2117–2134.

Ballesteros J.A., Sharma P.P. 2019. A Critical Appraisal of the Placement of Xiphosura (Chelicerata) with Account of Known Sources of Phylogenetic Error. Syst. Biol. 68:896–917.

Bailey T.L., Johnson J., Grant C.E., Noble W.S. 2015. The MEME Suite. Nucleic Acids Res 43:W39–W49.

Barchan D., Kachalsky S., Neumann D., Vogel Z., Ovadia M., Kochva E., Fuchs S. 1992. How the mongoose can fight the snake: The binding site of the mongoose acetylcholine receptor. Proc. Natl. Acad. Sci. USA 89:7717–7721.

Borges A., Graham M.R. 2014. Phylogenetics of Scorpions of Medical Importance. In: Venom Genomics and Proteomics (ed: Gopalakrishnakone P, Calvete JJ). Springer Press, Dordrecht, Germany, pp 81–104.

Cao Z., Yu Y., Wu Y., Hao P., Di Z., He Y., Chen Z., Yang W., Shen Z., He X., Sheng J., Xu X., Pan B., Feng J., Yang X., Hong W., Zhao W., Li Z., Huang K., Li T., Kong Y., Liu H., Jiang D., Zhang B., Hu J., Hu Y., Wang B., Dai J., Yuan B., Feng Y., Huang W., Xing X., Zhao G., Li X., Li Y., Li W. 2013. The genome of *Mesobuthus martensii* reveals a unique adaptation model of arthropods. Nat. Commun. 4:2602.

Casewell N.R., Wüster W., Vonk F.J., Harrison R.A., Fry B.G. 2013. Complex cocktails: the evolutionary novelty of venoms. Trends Ecol. Evol. 28:219–229.

Díaz-Perlas C., Varese M., Guardiola S., García J., Sánchez-Navarro M., Giralt E., Teixidó M. 2018. From venoms to BBB-shuttles. MiniCTX3: a molecular vector derived from scorpion venom. Chem. Commun. 54:12738–12741.

Drabeck D.H., Dean A.M., Jansa S.A. 2015. Why the honey badger don’t care: Convergent evolution of venom-targeted nicotinic acetylcholine receptors in mammals that survive venomous snake bites. Toxicon 99:68–72.

dos Reis M., Yang Z. 2019. Bayesian Molecular Clock Dating Using Genome-Scale Datasets. In: Anisimova M. (eds) Evolutionary Genomics. Methods in Molecular Biology, vol 1910. Humana, New York, NY, pp. 309–330.

Fet V., Gantenbein B., Gromov A.V., Lowe G., Lourenço W.R. 2003. The first molecular phylogeny of Buthidae (Scorpiones). Euscorpius 4:1–10.

Fet V., Soleglad M, Lowe G. 2005. A new trichobothrial character for the high-level systematics of Buthoidea (Scorpiones: Buthida). Euscorpius 23:1–40.

Frickey T., Lupas A. 2004. CLANS: a Java application for visualizing protein families based on pairwise similarity. Bioinformatics 20:3702–3704.

He Y., Zhao R., Di Z., Li Z., Xu X., Hong W., Wu Y., Zhao H., Li W., Cao Z. 2013. Molecular diversity of Chaerilidae venom peptides reveals the dynamic evolution of scorpion venom components from Buthidae to non-Buthidae. J. Proteomics 89:1–14.

Herzig V., Sunagar K., Wilson D.T.R., Pineda S.S., Israel M.R., Dutertre S., McFarland B.S., Undheim E.A.B., Hodgson W.C., Alewood P.F., Lewis R.J., Bosmans F., Vetter I., King G.F., Fry B.G. 2020. Australian funnel-web spiders evolved human-lethal δ-hexatoxins for defense against vertebrate predators. Proc. Natl. Acad. Sci. USA 117: 24920–24928.

Holding M.L., Drabeck D.H., Jansa S.A., Gibbs H.L. 2016. Venom Resistance as a Model for Understanding the Molecular Basis of Complex Coevolutionary Adaptations. Integr. Comp. Biol. 56:1032–1043.

Hopp B.H., Arvinson R.S., Adams M.E., Razak K.A. 2017. Arizona bark scorpion venom resistance in the pallid bat, *Antrozous pallidus*. PLoS ONE 12:e0183215.

King G.F., Hardy M.C. 2013. Spider-Venom Peptides: Structure, Pharmacology, and Potential for Control of Insect Pests. Ann. Rev. Entomol. 58:475–96.

Lartillot N., Philippe H. 2004. A Bayesian mixture model for across-site heterogeneities in the amino-acid replacement process. Mol. Biol. Evol. 21:1095–1109.

Mirarab S., Warnow T. 2015. ASTRAL-II: coalescent-based species tree estimation with many hundreds of taxa and thousands of genes. Bioinformatics 31:i44–i52.

Mistry J., Finn R.D., Eddy S.R., Bateman A., Punta M. 2013. Challenges in Homology Search: HMMER3 and Convergent Evolution of Coiled-Coil Regions. Nucleic Acids Res. 41:e121.

Niermann C., Tate T.G., Suto A.L., Barajas R., While H.A., Guswiler O.D., Secor S.M., Rowe A.H., Rowe M.P. 2020 Defensive Venoms: Is Pain Sufficient for Predator Deterrence? Toxins 12:260.

Nguyen L.T., Schmidt H.A., Haeseler A., Minh B.Q. 2014. IQ-TREE: a fast and effective stochastic algorithm for estimating maximum-likelihood phylogenies. Mol. Biol. Evol. 32:268–274.

Ojanguren-Affilastro A.A., Adilardi R.S., Mattoni C.I., Ramírez M.J., Ceccarelli F.S. 2017. Dated phylogenetic studies of the southernmost American buthids (Scorpiones; Buthidae). Mol. Phylogenet. Evol. 110:39–49.

Puttick M.N. 2019. MCMCtreeR: functions to prepare MCMCtree analyses and visualize posterior ages on trees. Bioinformatics 35(24):5321–5322.

Rapôso C. 2017 Scorpion and spider venoms in cancer treatment: state of the art, challenges, and perspectives. J. Clin. Transl. Res. 3:233–249.

Rowe A.H., Xiao Y., Rowe M.P., Cummins T.R., Zakon H.H. 2013. Voltage-gated sodium channel in grasshopper mice defends against bark scorpion toxin. Science 342:441–446.

Santibáñez-López C.E., Kriebel R., Ballesteros J.A., Rush N., Witter Z., Williams J., Janies D.A., Sharma P.P. 2018. Integration of phylogenomics and molecular modeling reveals lineage-specific diversification of toxins in scorpions. PeerJ 6:e5902–5923.

Santibáñez-López C.E., González-Santillán E., Monod L., Sharma P.P. 2019a. Phylogenomics facilitates stable scorpion systematics: Reassessing the relationships of Vaejovidae and a new higher-level classification of Scorpiones (Arachnida). Mol. Phylogenet. Evol. 135:22–30.

Santibáñez-López C.E., Graham M.R., Sharma P.P., Ortiz E., Possani L.D. 2019b. Hadrurid Scorpion Toxins: Evolutionary Conservation and Selective Pressures. Toxins 11:637.

Santibáñez-López C.E., Ojanguren-Affilastro A.A., Sharma P.P. 2020. Another one bites the dust: taxonomic sampling of a key genus in phylogenomic datasets reveals more non-monophyletic groups in traditional scorpion classification. Invertebr. Syst. 34:133–143.

Santos M.S., Silva C.G.L., Neto B.S., Júnior C.R.P., Lopes V.H.G., Júnior A.G.T., Bezerra D.A., Luna J.V.C.P., Cordeiro J.B., Júnior J.G., Lima M.A.P. 2016. Clinical and epidemiological aspects of scorpionism in the world: a systematic review. Wild Environ. Med. 27:504–518.

Schwager E.E., Sharma P.P., Clarke T., Leite D.J., Wierschin T., Pechmann M., Akiyama-Oda Y., Esposito L., Bechsgaard J., Bilde T., Buffry A.D., Chao H., Dinh H., Doddapaneni H., Dungan S., Eibner C., Extavour C.G., Funch P., Garb J., Gonzalez L.B., Gonzalez V.L., Griffiths-Jones, S., Han Y., Hayashi C., Hilbrant M., Hughes D.S.T., Janssen R., Lee S.L., Maeso I., Murali S.C., Muzny D.M., Nunes da Fonseca R., Paese C.L.B., Qu J., Roshaugen M., Schomburg C., Schönauer A., Stollewerk A., Torres-Oliva M., Turetzek N., Vanthournout B., Werren J.H., Wolf C., Worley K.C., Bucher G., Gibbs R.A., Coddington J., Oda H., Stanke M., Ayoub N.A., Prpic N.M., Flot J.F., Posnien N., Richards S., McGregor A.P. 2017. The house spider genome reveals an ancient whole-genome duplication during arachnid evolution. BMC Biol. 15:62.

Sharma P.P., Fernández R., Esposito L.A., González-Santillán E., Monod L. 2015. Phylogenomic resolution of scorpions reveals multilevel discordance with morphological phylogenetic signal. Proc. R. Soc. B 282:20142953.

Sharma P.P., Baker C.M., Cosgrove J.G., Johnson J.E., Oberski J.T., Raven R.J., Harvey M.S., Boyer S.L., Giribet G. 2018. A revised dated phylogeny of scorpions: Phylogenomic support for ancient divergence of the temperate Gondwanan family Bothriuridae. Mol. Phylogenet. Evol. 122:37–45.

Sissom W.D. 1990. Sytematics, Biogeography, and Paleontology. In: The Biology of Scorpions (ed: Polis GA). Stanford University Press, Stanford, CA, US, pp 31–80.

Suranse V., Sawant N.S., Paripatydar S.V., Krutha K., Paingankar M.S., Padhye A.D., Bastawade D.B., Dahanukar N. 2017. First molecular phylogeny of scorpions of the family Buthidae from India. Mitochondrial DNA A 28:606–611.

Tropea G., Onnis C. 2020. A remarkable discovery of a new scorpion genus and species from Sardinia (Scorpiones: Chactoidea: Belisariidae). Riv. Aracnol. Ital. 26:3–25.

Upham N.S., Esselstyn J.A., Jetz W. 2019. Inferring the mammal tree: Species-level sets of phylogenies for questions in ecology, evolution, and conservation. PLoS Biol. 17: e3000494.

Veiseh M., Gabikian P., Bahrami S.B., Veiseh O., Zhang M., Hackman R.C., Ravanpay A.C., Stroud M.R., Kusama Y., Hansen S.J., Kwok D., Munoz N.M., Sze R.W., Grady W.M., Greenberg N.M., Ellebogen R.G., Olson J.M. 2007. Tumor Paint: A Chlorotoxin:Cy5.5 Bioconjugate for Intraoperative Visualization of Cancer Foci. Cancer Res. 67:6882–6888.

Yang Z. 2007. PAML 4: phylogenetic analysis by maximum likelihood. Mol. Biol. Evol. 24:1586–1591.

