## Supplemental Material for "Phylogenomics of scorpions reveal a co-diversification of scorpion mammalian predators and mammal-specific sodium channel toxins"

Carlos E. Santibáñez-López

**This PDF file includes:**

Supplementary text

Supplementary References

Supplementary Figures S1 to S17

### Supplementary Text

#### *Taxon sampling*

Sequences from 100 species of Scorpiones were used for this study, from which 55 were sequenced previously by our team and 42 were newly sequenced for this study. In addition, 20 sequences of outgroup species representing seven chelicerate orders were included totaling 120 species. A full list of sequences with taxonomic information, sources of previously published sequenced, and GenBank accession numbers are provided in Supplementary Tables S1-3.

#### *RNA extraction and sequencing*

All specimens were either collected, or from captive bred colonies (Table S2). All scorpions were dissected into RNAlater solution (Ambion, Foster City, CA, USA). From all specimens, the brain, legs and telson were dissected for sequencing as in our previous publications (Santibáñez-López et al. 2019). Total RNA extraction, mRNA isolation and cDNA library construction were performed as in our previous protocols (Sharma et al. 2014; 2015; Santibáñez-López et al. 2018a; 2018b). RNA was extracted using the Trizol Trireagent system (Ambion Life Technologies, Waltham, MA, USA) Library preparation and stranded mRNA sequencing followed protocols from the Biotechnology Center at the University of Wisconsin-Madison or constructed in the Apollo 324 automated system using the PrepX mRNA kit (IntegenX, Pleasanton, CA, USA), with samples marked with unique indices to enable multiplexing. Samples were run using the Illumina HiSeq 2500 platform with paired-end reads of 100 or 150 bp at the FAS Center for Systems Biology at Harvard University, and using an Illumina HiSeq2500 High Throughput platform with paired-end reads of 125 bp at the Biotechnology Center at the University of Wisconsin-Madison.

*De novo transcriptome assemblies and orthology inference*

The 42 new transcriptomes were assembled using Trinity v. 2.5 (Grabherr et al. 2011), removing the adaptors with Trimmomatic v. 0.36 (Bolger et al. 2014) and assessing the quality of cleaned raw reads with FastQC v. 0.11.5 (Andrews 2010). Protein coding regions within the assembled transcripts were identified using TransDecoder v. 5.3.0 (Haas et al. 2013). De novo phylogenetic orthology resulted prohibitive for a dataset of this size. To overcome this limitation, we leveraged on the 3564 of orthologous computed from 53 chelicerates and outgroups as described in (Ballesteros and Sharma 2019). The untrimmed alignment of each pre-computed orthogroup was used to produce a hidden Markov profile using hmmrbuild from the hmmr package v.3.2.1(Finn et al. 2011), in doing so, the variation present in each group is parameterized and the number of searches is reduced to the number of profiles (genes). Each proteome/transcriptome of the species of interest was then searched (hmmrseacrh) for matches against the collection of profiles with an expectation threshold of $e < 10^{-20}$; for cases with more than one hits per locus, the sequence with the best score was selected, and the corresponding sequence appended to the locus FASTA file aggregating the putative orthologous found in each species.

These putative orthogroups was further refined by comparing each of its sequences against the proteome of the fruit-fly *Drosophila melanogaster*, identifying the most common "best hit" fruit-fly gene and removing from the orthogroup all the sequences pointing to a different best-hit with *D. melanogaster*. Notice that this filter does not require that *D. melanogaster* as member of the group.

##### *Phylogenetic methods*

From this collection of orthologs, gene trees were estimated using IQTREE v. 1.6. (Nguyen et al. 2014) implementing the best-fitting amino acid substitution model (-m MFP). Subsequently, these orthologs were subject to a relaxed occupancy filtering and finally, each gene tree was examined for paralogy and either removed or the whole locus discarded.

Three matrices were assembled with a minimal taxon occupancy threshold. For Matrix 1, we favored the presence of at least 115 species per ortholog; for matrix 2 the presence of at least 109 species and for matrix 3, the presence of at least 103 species. Matrix 1 consisted of 192 loci and 53,333 sites (>95% complete), matrix 2 consisted of 424 loci and 114,188 sites (>90% complete), and matrix 3 consisted of 660 loci and 185,631 sites (>85% complete). Phylogenetic inference of these concatenated matrices was computed with IQ-TREE implementing the best-fitting amino acid substitution per partition as selected in our gene tree estimation (-spp). Species tree estimation of the constituent orthologs of the three matrices were generated using the collection of ML gene trees and computed with ASTRAL v.3 (Mirarab and Warnow 2015). Lastly, while Bayesian inference analysis was explored for Matrix 3 (>95 complete) using Phylobayes-mpi v 1.7 (Lartillot et al. 2013) under a CAT + GTR + $\Gamma_4$ model failed to converge after over 5 months of computation time on four independent runs.

To further explore the generic grouping previously defined on the basis of trichobothria position on the surface of the pedipalps (*sensu* Fet et al. 2005), we searched GenBank for molecular data available for other scorpion genera/species not sequenced in this study (Supplementary Table S4). We retrieved eight scorpion species RNAseq data, one species EST datasets, and one genome [*Mesobuthus martensii* Karsch, (1879)]. The orthology inference of the 10 species was conducted as previously stated, using the collection of orthologs of our Matrix 2 (henceforth M2.plus).

In addition to this, sanger-sequences for 10 species (favoring buthid genera for which RNAseq is not available yet), such as the mitochondrial Cytochrome subunit I (COI) and 16s; and the nuclear ribosomal subunit 18s was retrieved from GenBank. Sequences of these loci were also included for the rest of the species when available. Preliminary ML phylogenetic analyses were conducted using these sequences with IQ-TREE implementing the best-fitting nucleotide substitution model with ModelFinder (Kalyaanamoorthy et al. 2017). Using the resulting topologies, we selected those sequences that recovered the correct species phylogenetic position, and subsequently added to our M2.plus matrix (henceforth, M2.plus.sanger). Phylogenetic inference of M2.plus.sanger was computed with IQ-TREE as mentioned previously (-spp).

*Divergence time estimation*

It is well known that performing divergence time estimation with large, modern datasets is notoriously computationally intensive. Thus, Matrix 1 was selected to estimate dates of divergence based on its composition (high number of loci and lower values of missing data) with MCMCTree and codeml from the PAML software package v 4.8 (Yang 2007) using the approximate likelihood method (dos Reis and Yang 2011). The input tree was the best ML topology from the analysis of Matrix 1 and was time-calibrated using 15 calibration points. These points were selected based on several references and are listed in the Supplementary Table S5. Nine non-scorpion calibration points were introduced as minimum bounds of uniform prior, the six scorpion calibration points with minimum and maximum bounds, and with the maximum root age as a hard maximum bound. A conservative age constraint on the basal diversification of Arachnida was applied (see *Fossil calibrations*, below).

As suggested by other studies (i.e. Kawahara et al. 2019), the Dirichlet-Gamma density (dos Reis et al. 2014) was used to set the prior on the molecular rate with the default parameters ($\alpha_\mu = 2$, $\beta_\mu = 6$, $\alpha = 1$). Similarly, the default parameters for the birth-death process were kept ( $\lambda = 1$, $p = 0.1$). Independent-rates and the correlated rates models were selected to perform the divergence time estimation. Hessian matrices were calculated with *codeml* using empirical base frequencies and the LG substitution model with 5 rate categories. A burn-in of 25,000, sample frequency of 100 and a sample size of 25,000 (resulting in a 2.5 million generation) for each of the two clock models were settled. Each analysis was repeated four times, with the convergence subsequently evaluated in Tracer v. 1.7.1 (Rambaut et al. 2018) and by plotting the node ages against each other in R (data not shown). Lastly, the four chains were merged to reach an effective sample size (ESS) greater than 200 and produced a summary tree. Resulting topology was plotted using the function *MCMC.tree.plot* from the R package MCMCtreeR (Puttick 2019) and edited in Adobe Illustrator ®.

The posterior distribution of node ages of Didelphimorphia, Eulipotyphia, Chiroptera, Carnivora and Rodentia were generated from the 95% credibility intervals from (Upham et al. 2019); filled with random ages from a normal distribution and a standard deviation of 0.1 plotted using the function *node.posteriors* from MCMCtreeR. On the other hand, the posterior distribution of nodes ages of the selected clades of scorpions were retrieved from our results.

##### *Fossil calibrations*

The oldest unequivocal fossil scorpion, *Parioscorpio venator*, was recently discovered and dates to the Silurian (436.5-437.5 Mya; Wendruff et al. 2020), slightly pushing back the age of the oldest stem-group Scorpiones as inferred from fossils like *Dolichophonus loudenensis* and *Eramoscropius brucensis*; the oldest crown group scorpion (Orthosterni) is Carboniferous in age (*Protoischnurus axelrodorum*) (reviewed by Wolfe et al. 2016). We constrained the stem age of scorpions (divergence from outgroup taxa) using a soft minimum of 435.15 Mya and a soft maximum of 514 Mya, with the upper bound following an established calibration strategy for arthropods (Wolfe et al. 2016). We constrained crown Orthosterni with a soft maximum age of 313.7 Mya and a soft minimum age of 112.6 Mya, based on the age of *Compsoscorpius buthiformis*, a fossil Scorpionoidea (Poiton et al. 2012). Given the clear affinity of *Protoischnurus axelrodorum* as a lineage within Iurida (Menton 2007), Iurida was also constrained using soft bounds of 112.6 to 313.7 Mya. Chaerilidae was constrained using a soft minimum age of 98.17 Mya, based on the age of *Electrochaerilus buckleyi* (Santiago-Blay et al. 2004a) and a soft maximum age of 313.7 Mya. We constrained Buthoidea using a soft minimum of 49.26 Mya, based on the age of *Uintascorpio halandrasi* (Perry 1995, Santiago-Blay et al. 2004b) and a soft maximum age of 313.7 Mya, given the sister group relationship of Buthida and Iurida. Finally, the superfamily Scorpionoidea was constrained with a soft minimum of 112.6 Mya, based on the age of *Compsoscorpius buthiformis*, and a soft maximum of 313.7 Mya. Further discussion of these fossils is provided in a recent review (Howard et al. 2019).

Ten additional outgroup calibrations were deployed for the remaining chelicerate orders, using only soft minimum calibrations. We constrained the crown group of Opiliones using the soft minimum age of 411 Mya, based on the harvestman fossil *Eophalangium sheari* (Garwood et al. 2014), the divergence of Eupnoi harvestmen with a soft minimum age of 305 Mya, based on the age of *Macrogyion cronus* (Garwood et al. 2011); and the divergence of Dyspnoi harvestmen with a soft minimum age of 305 Mya, based on the age of *Ameticos scolos* (Garwood et al. 2011). The age of Araneae was calibrated with a soft minimum age of 386 Mya, based on the fossil *Attercopus fimbriunguis* (Selden et al. 2008, Huang et al. 2018). The stem-group age of Mesothelae was calibrated with a soft minimum age of 305 Mya, based on the unambigious mesothele fossil *Palaeothele montceauensis* (Selden 1996). The stem-group age of Amblypygi was bounded with a soft minimum age of 312 Mya based on the fossils *Graeophonus anglicus* and *G. carbonarius* (Dunlop et al. 2007). The crown group of Pseudoscorpiones was calibrated with a soft mimimum age of 392 Mya, based on the age of *Dracochela deprehendor* (Schawaller et al. 1991). Finally, the stem-group age of Xiphosura was constrained with a soft minimum age of 445 Mya, corresponding to the age of *Lunataspis aurora* (Rudkin et al. 2008).

*Gene tree analyses and molecular evolution of venom scorpion peptides*

Cysteine-stabilized $\alpha$-helix and $\beta$-sheet fold (CS$\alpha\beta$), disulphide-directed beta-hairpin (DDH) and Inhibitor cystine knot (ICK) homologs from scorpion venom were retrieved from our concatenated transcriptome libraries using a query files with known CS$\alpha\beta$ - ICK - DDH sequences from the venom of scorpions (i.e. sodium channel toxins, potassium channel toxins, chloride channel toxins, scorpines, Kunitz-type inhibitors, and calcins). Matching sequences with low e-values (< 1 x $10^{-15}$) were selected. Subsequently, these selected sequences were concatenated with 749 sequences from scorpion venom peptides retrieved from UniProt and InterPro (Supplementary Table S6).

Preliminary ML phylogenetic analyses were conducted using the entire dataset and IQ-TREE, with the resulting trees analyzed in search of orthologs. We removed sequences retrieved from our transcriptomes that were recovered outside clades with at least one known venom sequence. Our final dataset consisted of 1,353 CSαβ - ICK scorpion sequences and 41 DDH scorpion toxin sequences as outgroup. Clades were delimitated manually based on the presence of well-known scorpion toxins (Supplementary Table S6). Thus, the sodium channel toxins (NaTx) dataset consisted of 661 sequences, the potassium channel toxins (KTx) dataset of 550, the chloride channel toxins (ClTx) dataset of 56, and the calcin dataset of 74 sequences. Multiple sequence alignments for sequences from each clade using MAFFT and gene trees were generated using IQ-TREE. Our favored topology of NaTx (Figure 3B) was plotted and annotated with Iroki (Moore et al. 2020) and subsequently edited with Adobe Illustrator. The signal peptides, propeptides and mature peptides were predicted for the sequences in each clade using SpiderP from Archnoserver (Herzig et al 2010). To determine if convergent sites predates the origin of different clades of Mammal-specific toxins within NaTx, motifs from the mature peptide (the active component of the toxin) were generated using the Multiple Em for Motif Elicitation server (MEME 5.1.1 at http://meme-suite.org/tools/meme; Bailey et al. 2015).

Additionally, the mature peptides of the amino acid sequences from the two main clades recovered in the ML analysis of the NaTx (Aah2-like and Cn2-like) were used for CLANS cluster analysis (Frickey and Lupas 2004) with the default parameters as stated in the configuration file. Lastly, the Ancestral State Reconstruction (ASR) of the Aah2 and Cn2 clades were recovered using ProtASR2 (Arenas and Bastolla 2020) with default parameters, the pdb files 1ptx

(Aah2) and 1cn2 (Cn2), and the sequences from our consensus multiple alignment in Figure S16. Structure models for *B. tunetanus* insect-specific toxin (Q17254), and *L. hebraeus* (P0C5I5) were generated using the mature peptide in the SWISS-MODEL server (Biasini et al 2014).

#### Supplementary References


Andrews S. 2010. FastQC: a quality control tool for high throughput sequence data.

Arenas M., Bastolla U. 2020. ProtASR2: Ancestral reconstruction of protein sequences accounting for folding stability. Methods Ecol. Evol. 11: 248-257.

Bailey T.L., Johnson J., Grant C.E., Noble W.S. 2015. The MEME Suite. Nucleic Acids Res. 43(W1):W39–W49.

Ballesteros J.A., Sharma P.P. 2019. A Critical Appraisal of the Placement of Xiphosura (Chelicerata) with Account of Known Sources of Phylogenetic Error. Systematic Biology. 33:440–22.

Biasini M., Bienert S., Waterhouse A., Arnold K., Studer G., Schmidt T., Kiefer F., Cassarino T.G., Bertoni M., Bordoli L., Schwede T. 2014. SWISS-MODEL: modelling protein tertiary and quaternary structure using evolutionary information. Nucleic Acids Res. 42(W1): W252-258.

Bolger A.M., Lohse M., Usadel B. 2014. Trimmomatic: a flexible trimmer for Illumina sequence data. Bioinformatics. 30:2114–2120.

dos Reis M., Zhu T., Yang Z. 2014. The Impact of the Rate Prior on Bayesian Estimation of Divergence Times with Multiple Loci. Systematic Biology. 63:555–565.

dos Reis M.D., Yang Z. 2011. Approximate Likelihood Calculation on a Phylogeny for Bayesian Estimation of Divergence Times. Molecular Biology and Evolution. 28:2161–2172.

Dunlop J.A., Zhou G.R.S., Braddy S.J. 2007. The affinities of the Carboniferous whip spider *Graeophonus anglicus* Pocock, 1911 (Arachnida: Amblypygi). Earth Environ. Sci. Trans. R. Soc. Edinb. 98:165–178.

Fet V., Soleglad M.E., Lowe G. 2005. A new trichobotrhial character for the high-level systematics of Buthoidea (Scorpiones: Buthida). Euscorpius. 23:1–40.

Finn R.D., Clements J., Eddy S.R. 2011. HMMER web server: interactive sequence similarity searching. Nucleic Acids Res. 39:W29–W37.

Frickey T., Lupas A. 2004. CLANS: a Java application for visualizing protein families based on pairwise similarity. Bioinformatics 20(18):3702–3704.

Garwood R.J., Sharma P.P., Dunlop J.A., Giribet G. 2014. A Paleozoic stem group to mite harvestmen revealed through integration of phylogenetics and development. Curr. Biol. 24:1017–1023.

Garwood R.J, Dunlop J.A., Giribet G., Sutton M.D. 2011. Anatomically modern Carboniferous harvestmen demonstrate early cladogenesis and stasis in Opiliones. Nat. Commun. 2:444.

Grabherr M.G., Haas B.J., Yassour M., Levin J.Z., Thompson D.A., Amit I., Adiconis X., Fan L., Raychowdhury R., Zeng Q., Chen Z., Mauceli E.,

Hacohen N., Gnirke A., Rhind N., di Palma F., Birren B.W., Nusbaum C., Lindblad-Toh K., Friedman N., Regev A. 2011. Full-length transcriptome assembly from RNA-Seq data without a reference genome. Nat Biotechnol. 29:644–652.

Haas B.J., Papanicolaou A., Yassour M., Grabherr M., Blood P.D., Bowden J., Couger M.B., Eccles D., Li B., Lieber M., MacManes M.D., Ott M., Orvis J., Pochet N., Strozzi F., Weeks N., Westerman R., William T., Dewey C.N., Henschel R., LeDuc R.D., Friedman N., Regev A. 2013. De novo transcript sequence reconstruction from RNA-seq using the Trinity platform for reference generation and analysis. Nature Protocols. 8:1494–1512.

Herzig V., Wood D.L.A., Newell F., Chaumeil P.A., Kaas Q., Binford G.J., Nicholson G.M., Gorse D., King G.F. 2010. ArachnoServer 2.0, an updated online resource for spider toxin sequences and structures. Nucleic Acids Res. 39:D653–D657.

Howard R.J., Edgecombe G.D., Legg D.A., Pisani D., Lozano-Fernandez J. 2019. Exploring the evolution and terrestrialization of scorpions (Arachnida: Scorpiones) with rocks and clocks. Org. Div. Evol. 19:71–86.

Huang D., Hormiga G., Cai C., Su Y., Yin Z., Xia F., Giribet G. 2018. Origin of spiders and their spinning organs illuminated by mid-Cretaceous amber fossils. Nature Ecol. Evol. 2:623–627.

Kalyaanamoorthy S., Minh B.Q., Wong T.K.F., Haeseler von A., Jermiin L.S. 2017. ModelFinder: fast model selection for accurate phylogenetic estimates. Nat. Methods. 14:587–589.

Kawahara A.Y., Plotkin D., Espeland M., Meusemann K., Toussaint E.F.A., Donath A., Gimnich F., Frandsen P.B., Zwick A., Reis dos M., Barber J.R., Peters R.S., Liu S., Zhou X., Mayer C., Podsiadlowski L., Storer C., Yack J.E., Misof B., Breinholt J.W. 2019. Phylogenomics reveals the evolutionary timing and pattern of butterflies and moths. Proc. Natl. Acad. Sci. U.S.A. 116:22657–22663.

Lartillot N., Rodrigue N., Stubbs D., Richer J. 2013. PhyloBayes MPI: Phylogenetic Reconstruction with Infinite Mixtures of Profiles in a Parallel Environment. Systematic Biology. 62:611–615.

Menon F. 2007. Higher systematics of scorpions from the Crato Formation, Lower Cretaceous of Brazil. Palaeontology 50:185–195.

Mirarab S., Warnow T. 2015. ASTRAL-II: coalescent-based species tree estimation with many hundreds of taxa and thousands of genes. Bioinformatics. 31:44–52.

Moore R.M., Harrison A.O., McAllister S.M., Polson S.W., Wommack K.E. 2020. Iroki: automatic customization and visualization of phylogenetic trees. PeerJ 8:e8584.

Nguyen L.-T., Schmidt H.A., Haeseler von A., Minh B.Q. 2014. IQ-TREE: A Fast and Effective Stochastic Algorithm for Estimating Maximum-Likelihood Phylogenies. Molecular Biology and Evolution. 32:268–274.

Perry M.L. 1995. Preliminary description of a new fossil scorpion from the Middle Eocene Green River Formation, Rio Blanco County, Colorado. In: Dayvault RD, Averett WR (eds.), *The Green River Formation in Piceance Creek and*

*Eastern Uinta Basins Field Trip* (pp. 131–133). Grand Junction: Grand Junction Geological Society.

Pointon M.A., Chew D.M., Ovtcharova M., Sevastopulo G.D., Crowley Q.G. 2012. New high-precision U–Pb dates from western European Carboniferous tuffs; implications for time scale calibration, the periodicity of late Carboniferous cycles and stratigraphical correlation. J. Geol. Soc. Lond. 169:713–721.

Puttick M.N. 2019. MCMCtreeR: functions to prepare MCMCtree analyses and visualize posterior ages on trees. Bioinformatics. 35:5321–5322.

Rambaut A., Drummond A.J., Xie D., Baele G., Suchard M.A. 2018. Posterior Summarization in Bayesian Phylogenetics Using Tracer 1.7. Systematic Biology. 67:901–904.

Rudkin D.M., Young G.A., Nowlan G.S. 2008. The oldest horseshoe crab: a new xiphosurid from late Ordovician Konservat-Lagerstätten deposits, Manitoba, Canada. Palaeontology 51:1–9.

Santiago-Blay J.A., Fet V., Soleglad M.E., Anderson S.R. 2004a. A new genus and subfamily of scorpions from Cretaceous Burmese amber (Scorpiones: Chaerilidae). Rev. Ibér. Aracnol. 9:3–14.

Santiago-Blay J.A., Soleglad M.E., Fet V. 2004b. A redescription and family placement of *Uintascorpio* Perry, 1995 from the Parachute Creek Member of the Green River Formation (Middle Eocene) of Colorado, USA (Scorpiones: Buthidae). Rev. Ibér. Aracnol. 10:7–16.

Santibáñez-López C., Ontano A., Harvey M., Sharma P. 2018a. Transcriptomic Analysis of Pseudoscorpion Venom Reveals a Unique Cocktail Dominated by Enzymes and Protease Inhibitors. Toxins. 10:207–12.

Santibáñez-López C.E., González-Santillán E., Monod L., Sharma P.P. 2019. Phylogenomics facilitates stable scorpion systematics_ Reassessing the relationships of Vaejovidae and a new higher-level classification of Scorpiones (Arachnida). Molecular Phylogenetics and Evolution. 135:22–30.

Santibáñez-López C.E., Kriebel R., Ballesteros J.A., Rush N., Witter Z., Williams J., Janies D.A., Sharma P.P. 2018b. Integration of phylogenomics and molecular modeling reveals lineage-specific diversification of toxins in scorpions. PeerJ. 6:e5902–23.

Schawaller W., Shear W.A., Bonamo P.M. 1991. The first Paleozoic pseudoscorpions (Arachnida, Pseudoscorpionida). Amer. Mus. Novitates 3009:1–17.

Selden P.A. 1996 First fossil mesothele spider, from the Carboniferous of France. Rev. Suisse Zool. hors. série:585–596.

Selden P.A., Shear W.A., Sutton M.D. 2008. Fossil evidence for the origin of spider spinnerets, and a proposed arachnid order. Proc. Natl. Acad. Sci. USA 105:20781–20785.

Sharma P.P., Fernández R., Esposito L.A., González-Santillán E., Monod L. 2015. Phylogenomic resolution of scorpions reveals multilevel discordance with morphological phylogenetic signal. Proc. Biol. Sci. 282:20142953.

Sharma P.P., Kaluziak S.T., Pérez-Porro A.R., González V.L., Hormiga G., Wheeler W.C., Giribet G. 2014. Phylogenomic interrogation of arachnida reveals systemic conflicts in phylogenetic signal. Molecular Biology and Evolution. 31:2963–2984.

Upham N.S., Esselstyn J.A., Jetz W. 2019. Inferring the mammal tree: Species-level sets of phylogenies for questions in ecology, evolution, and conservation. PLoS Biol. 17:e3000494–44.

Wendruff A.J., Babcock L.E., Wirkner C.S., Kluessendorf J., Mikulic D.G. 2020. A Silurian ancestral scorpion with fossilised internal anatomy illustrating a pathway to arachnid terrestrialisation. Sci. Rep. 10:14.

Wolfe J.M., Daley A.C., Legg D.A., Edgecombe G.D. 2016 Fossil calibrations for the arthropod Tree of Life. Earth-Sci. Rev. 160:43–110.

Yang Z. 2007. PAML 4: Phylogenetic Analysis by Maximum Likelihood. Molecular Biology and Evolution. 24:1586–1591.

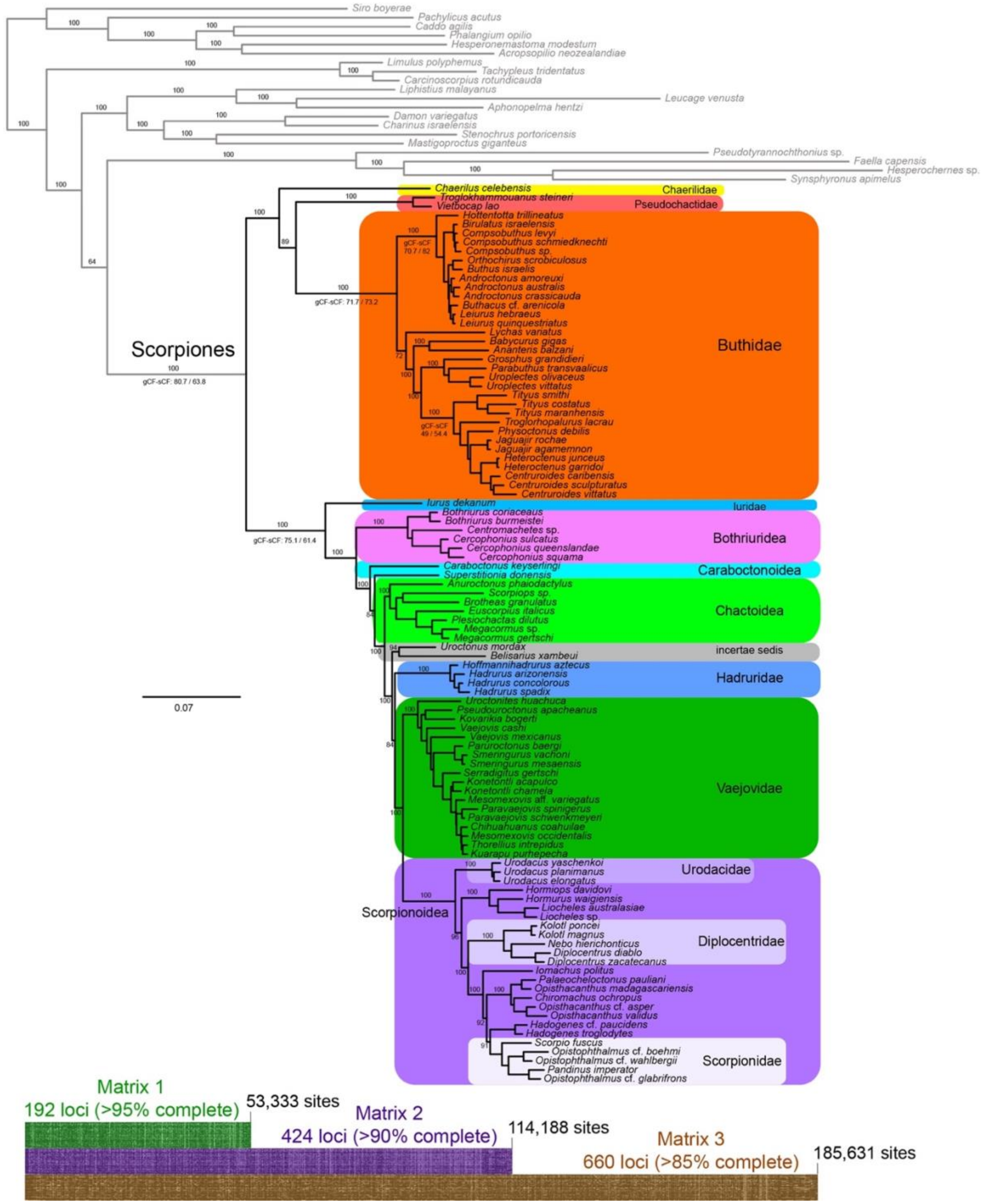


**Supplementary figure S1.** Maximum likelihood tree topology recovered from the analysis of 192 loci (Matrix 1). Numbers above nodes indicate bootstrap support values, below nodes indicate gene concordance factor (gCF) and the site concordance factor (sCF). Bottom panel shows an overview of the three matrices.

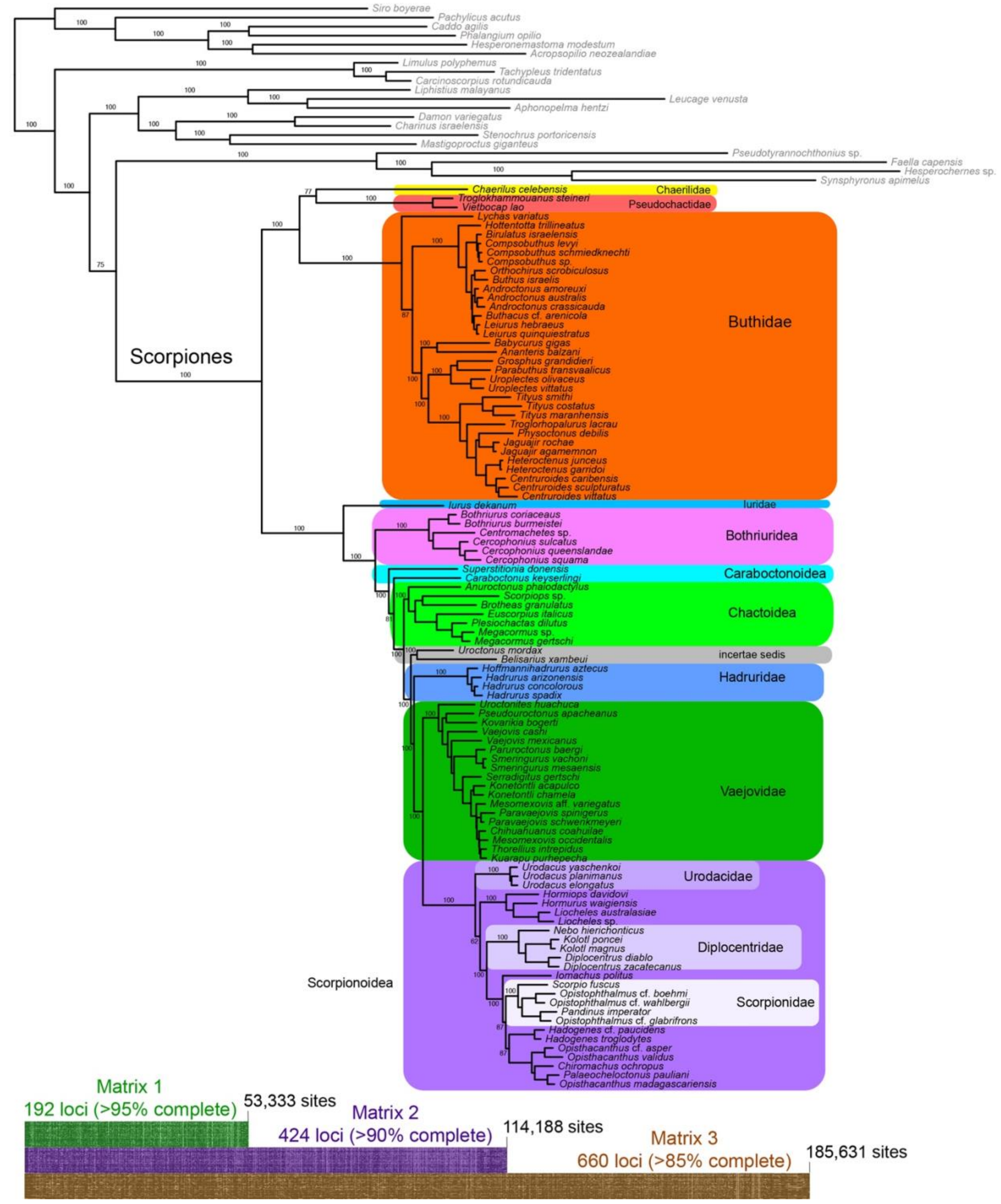


**Supplementary figure S2.** Maximum likelihood tree topology recovered from the analysis of 424 loci (Matrix 2). Numbers above nodes indicate bootstrap support values. Bottom panel shows an overview of the three matrices.

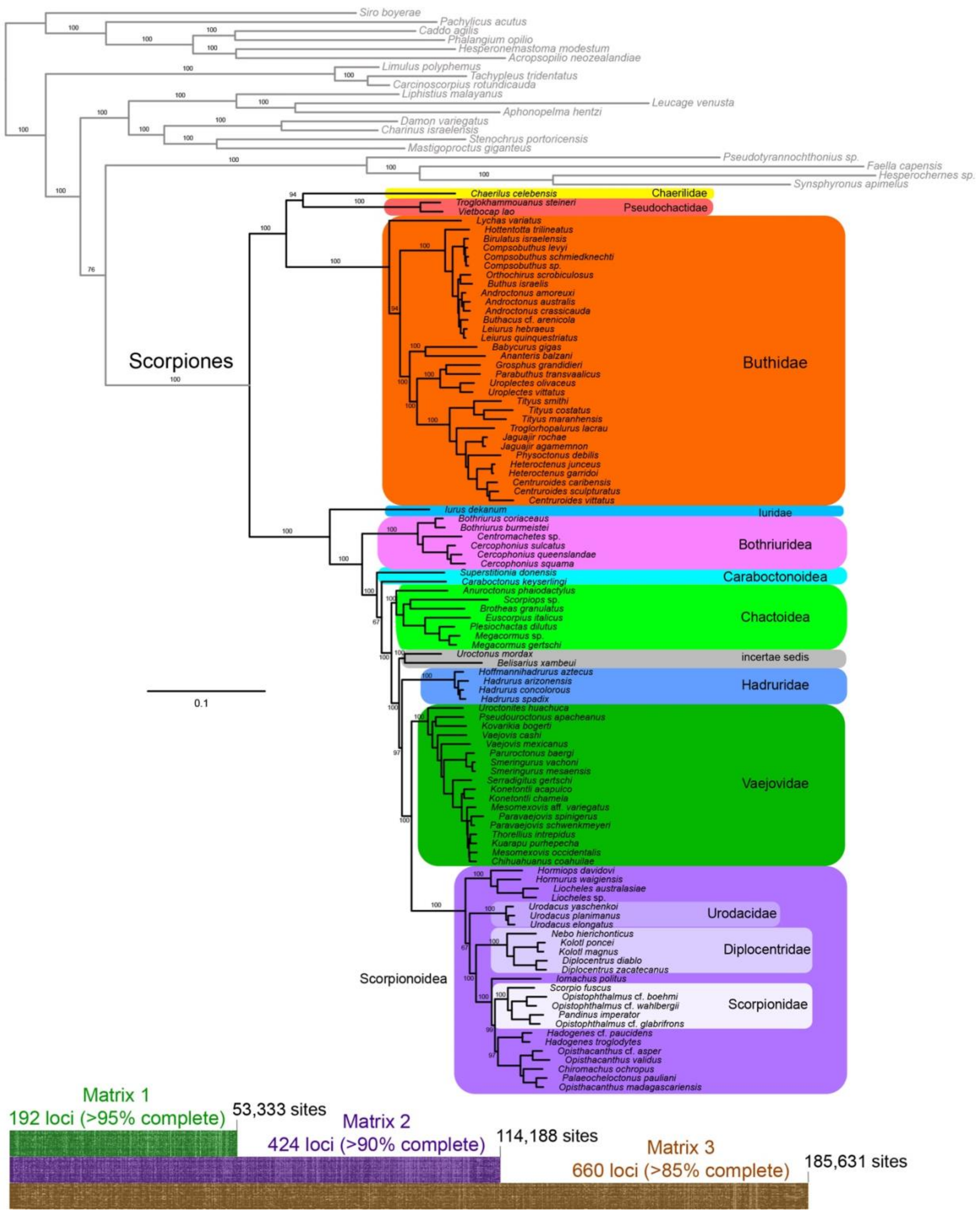


**Supplementary figure S3.** Maximum likelihood tree topology recovered from the analysis of 660 loci (Matrix 3). Numbers above nodes indicate bootstrap support values. Bottom panel shows an overview of the three matrices.

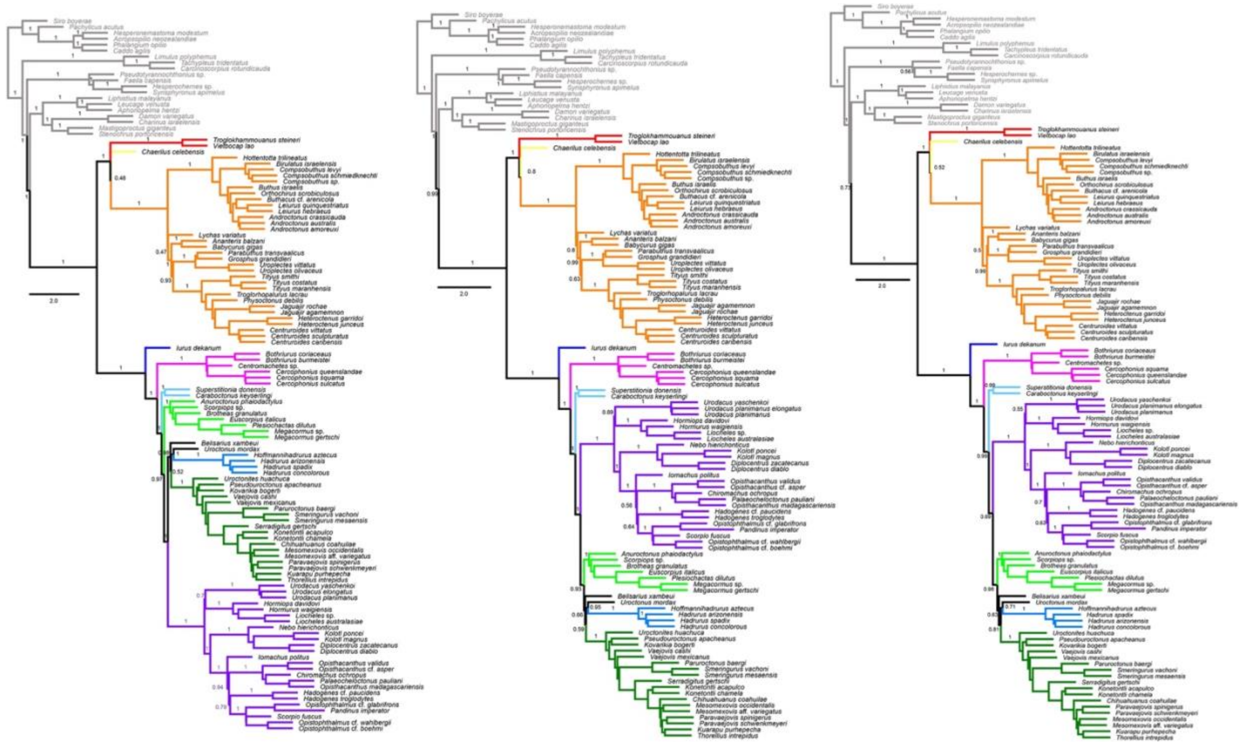

**Supplementary figure S4.** ASTRAL trees recovered from the 192- (left), 424- (center), and 660- (right) locus matrices. Numbers indicate branch support from local posterior probabilities.

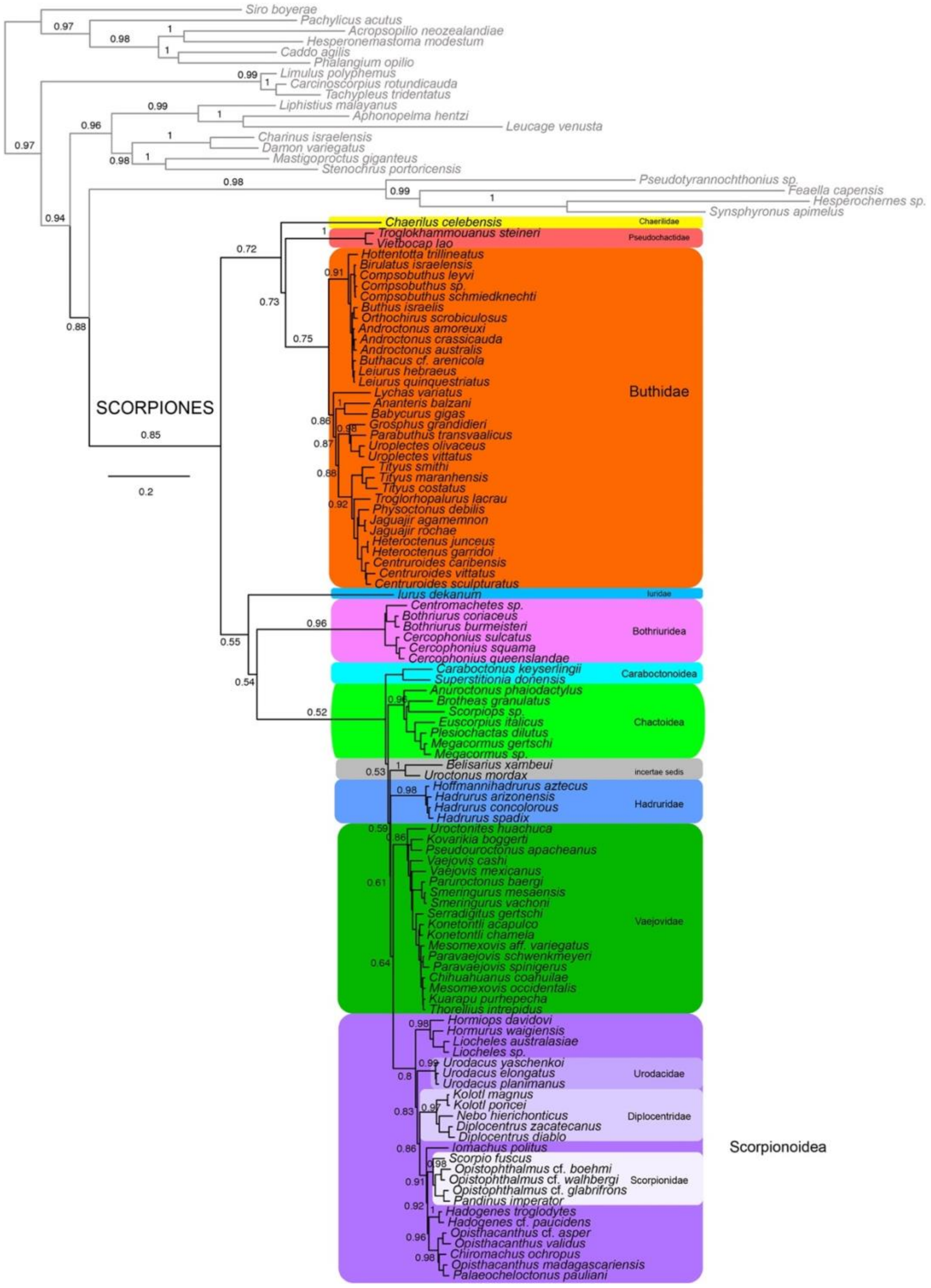


**Supplementary figure S5.** Bayesian inference tree topology recovered from the analysis of the 192-locus matrix with four independent chains under the CAT + GTR + $\Gamma_4$ model. Numbers on nodes indicate posterior probabilities.

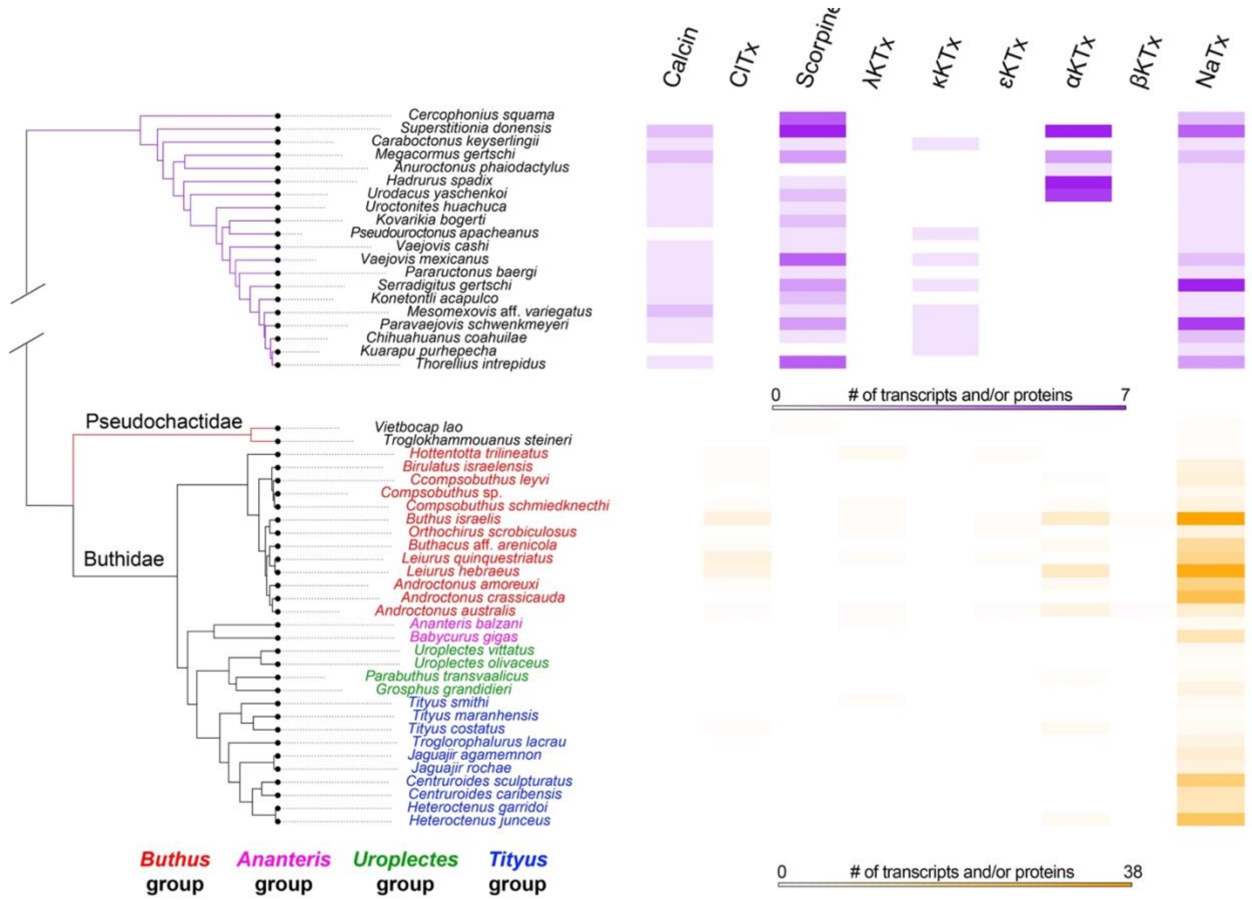


**Supplementary figure S6.** Summary of scorpion time tree showing the diversity of venom components in the different families and genera. Data were obtained from transcriptomic resources and databases (UniProt and InterPro).

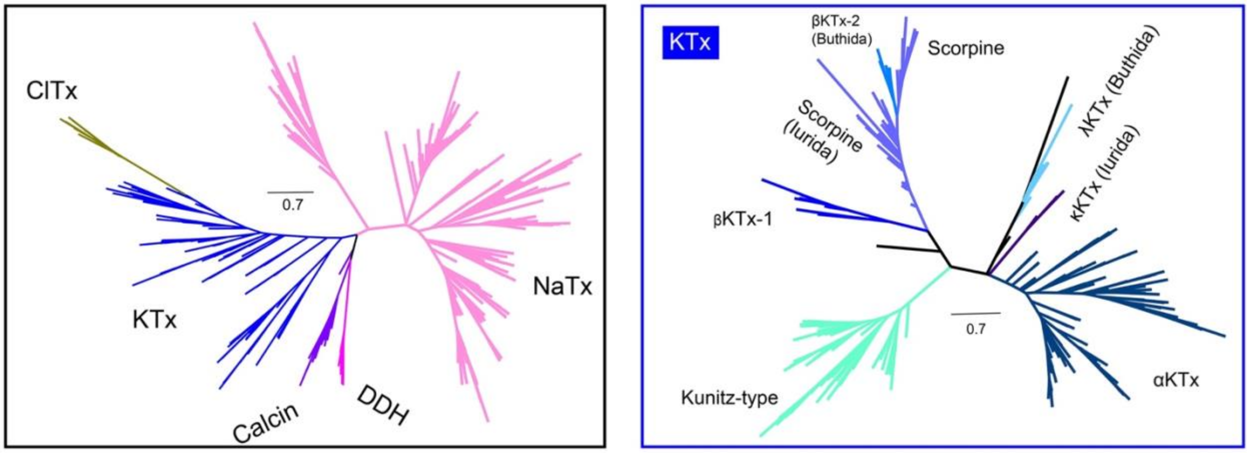


**Supplementary figure S7.** Maximum likelihood evolutionary tree for CSαβ-ICK gene families (left) and KTx gene families (right).

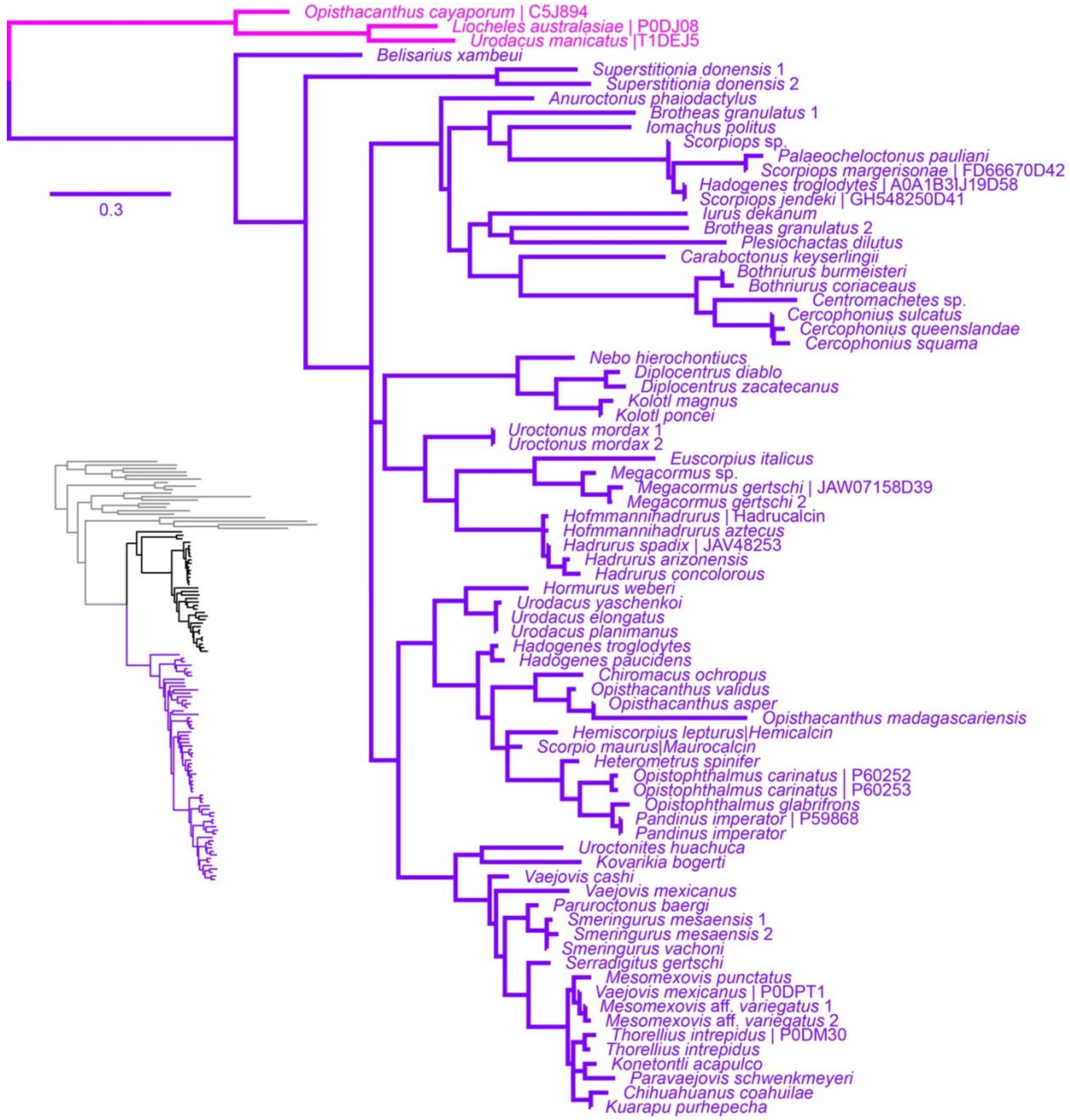


**Supplementary figure S8.** Maximum likelihood gene tree recovered from the analysis of 74 ryanodine receptor ligand peptides (calcins) and three disulphide-directed beta-hairpin (DDH) genes as outgroups. Calcins are ubiquitous in iurid scorpion venom as shown on the inset scorpion phylogeny.

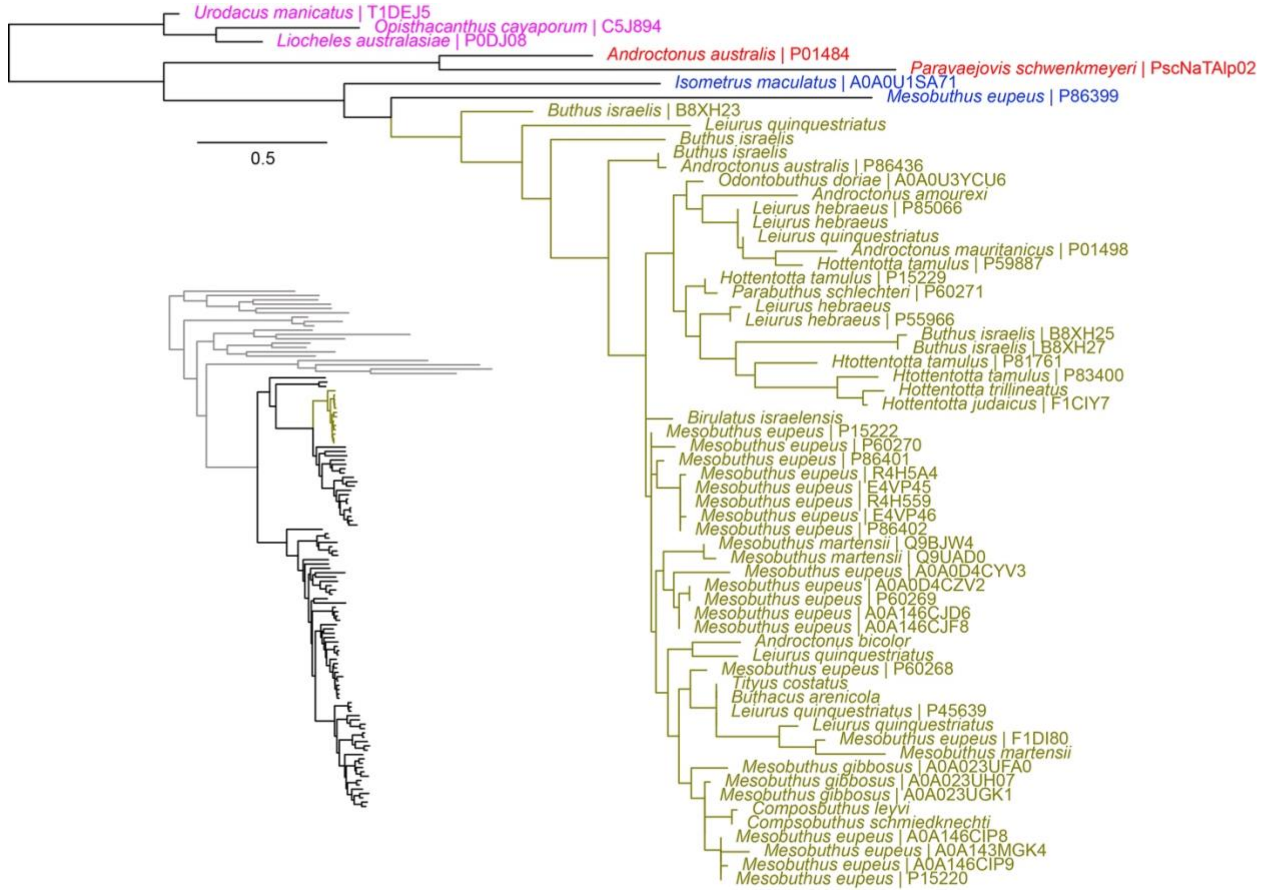


**Supplementary figure S9.** Maximum likelihood gene tree recovered from the analysis of 56 chloride channel toxin (ClTx) peptides and seven outgroups. ClTx are unique to buthid scorpions as shown on the inset scorpion phylogeny.

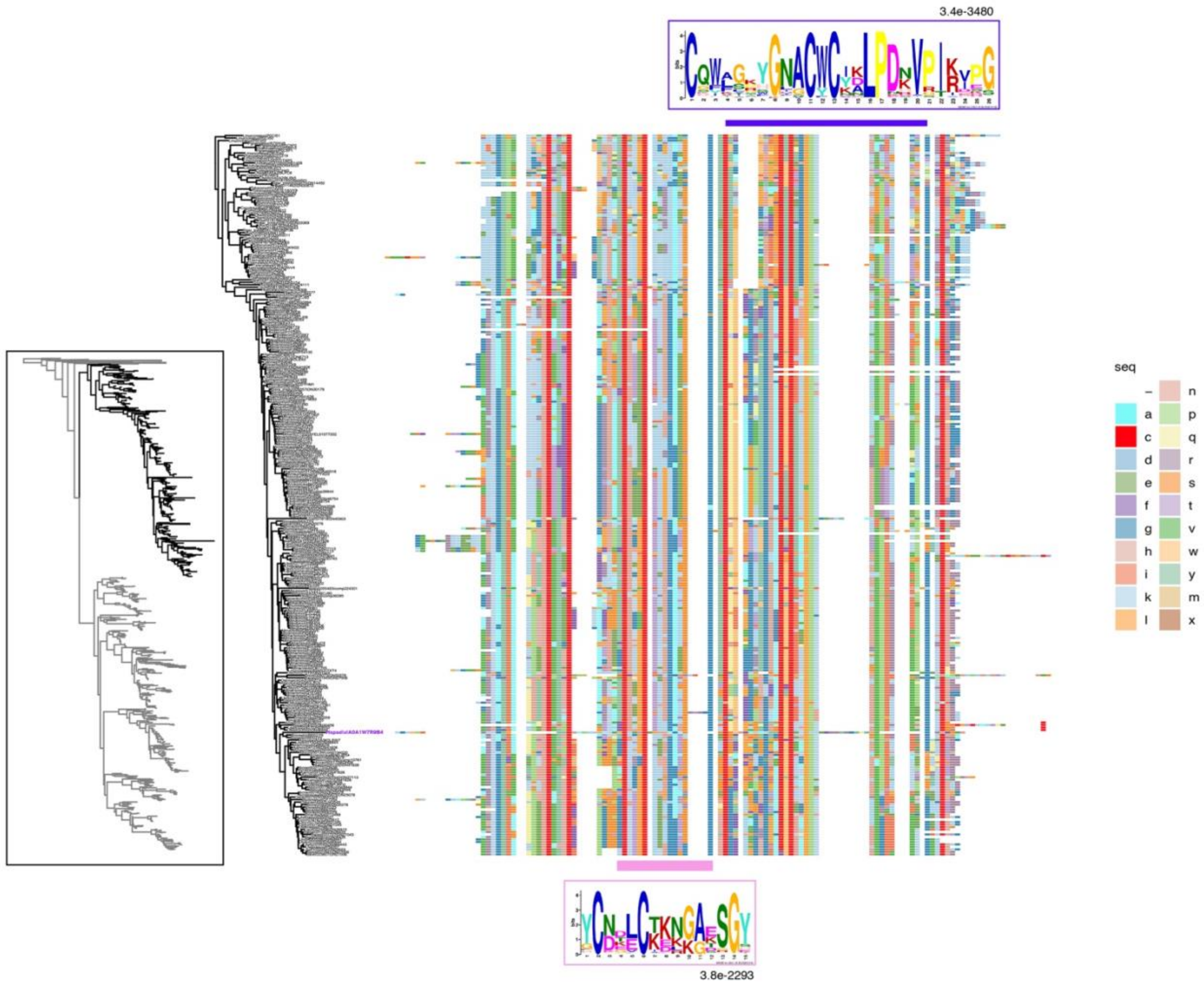


**Supplementary figure S10.** Maximum likelihood gene tree recovered from the analysis of 723 Sodium channel toxin (NaTx) peptides recovered from our transcriptomic analyses and databases (UniProt and InterPro), plus two outgroups (inset). Multiple sequence alignment color-coded for the 294 peptides from the Aah2-like clade (highlighted in black on inset phylogeny). Mature peptide repetitive motifs found using Multiple Em for Motif Elicitation (MEME) locations as shown by colored bars.

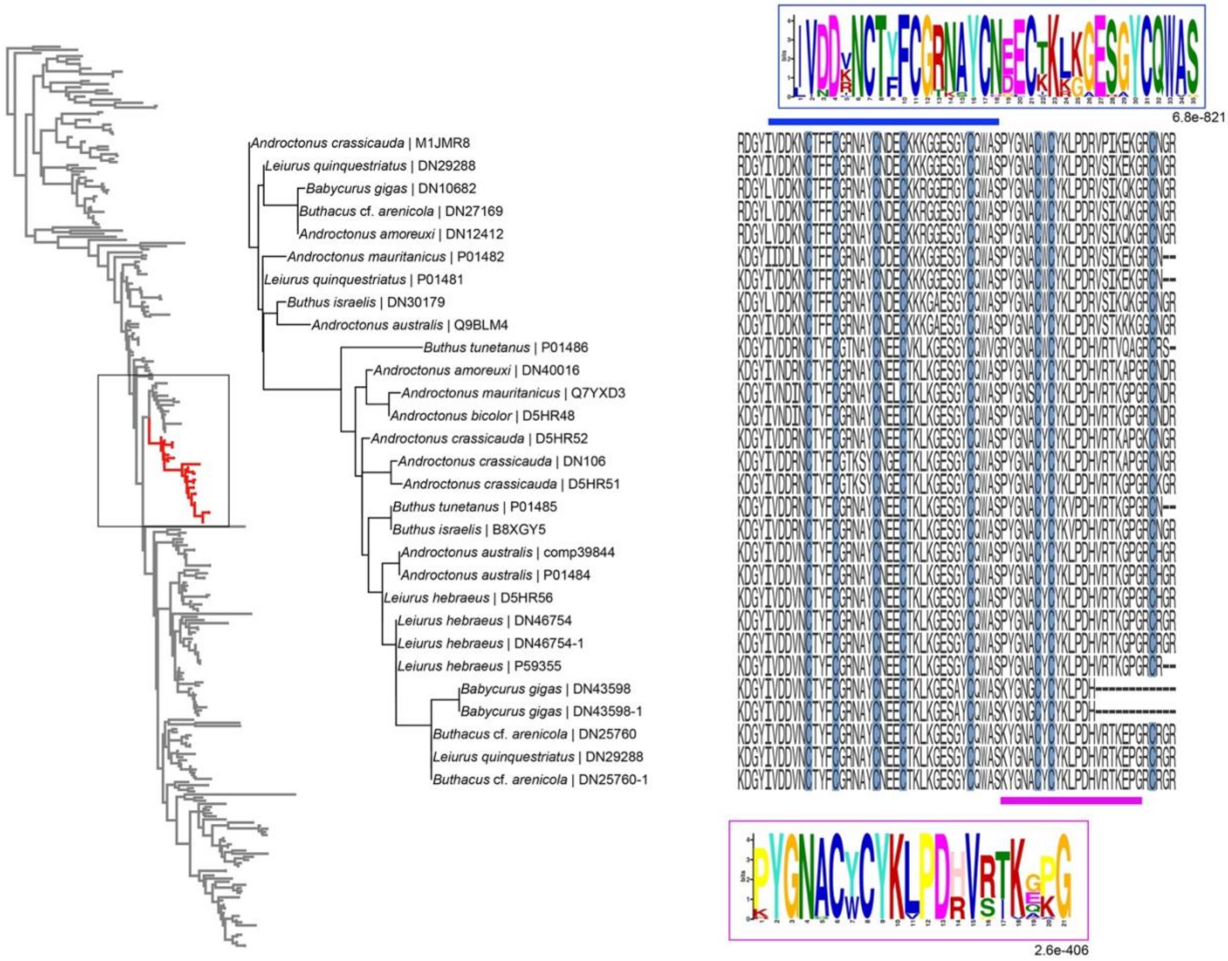


**Supplementary figure S11** Left: Pruned gene tree of the Aah2-like clade consisting of species from the Buthus groups. Two sequences from *Babycurus gigas* might be cross contamination from the Trinity assembly (see their high similarity to other sequences from toxic species). Right: Multiple sequence alignment of the mature peptide showing the repetitive motifs found using Multiple Em for Motif Elicitation (MEME).

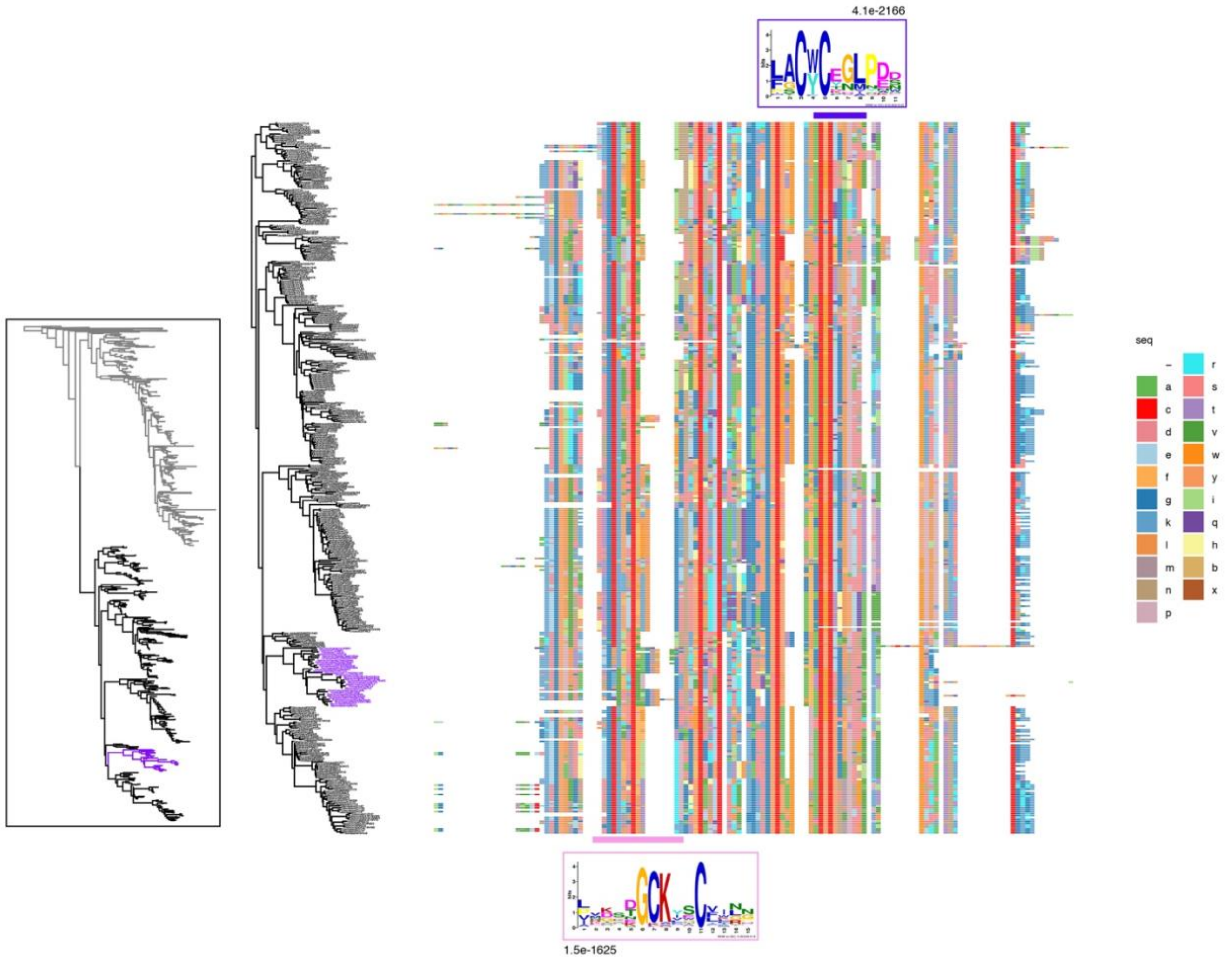


**Supplementary figure S12.** Maximum likelihood gene tree recovered from the analysis of 723 sodium channel toxin (NaTx) peptides recovered from our transcriptomic analyses and databases (UniProt and InterPro), plus two outgroups (inset). Multiple sequence alignment color-coded for the 376 peptides from the Cn2-like clade (highlighted in black on inset phylogeny). Mature peptide repetitive motifs found using Multiple Em for Motif Elicitation (MEME) locations as shown by colored bars. Highlighted in purple is the sole clade of iurid scorpion Cn2-like peptides.

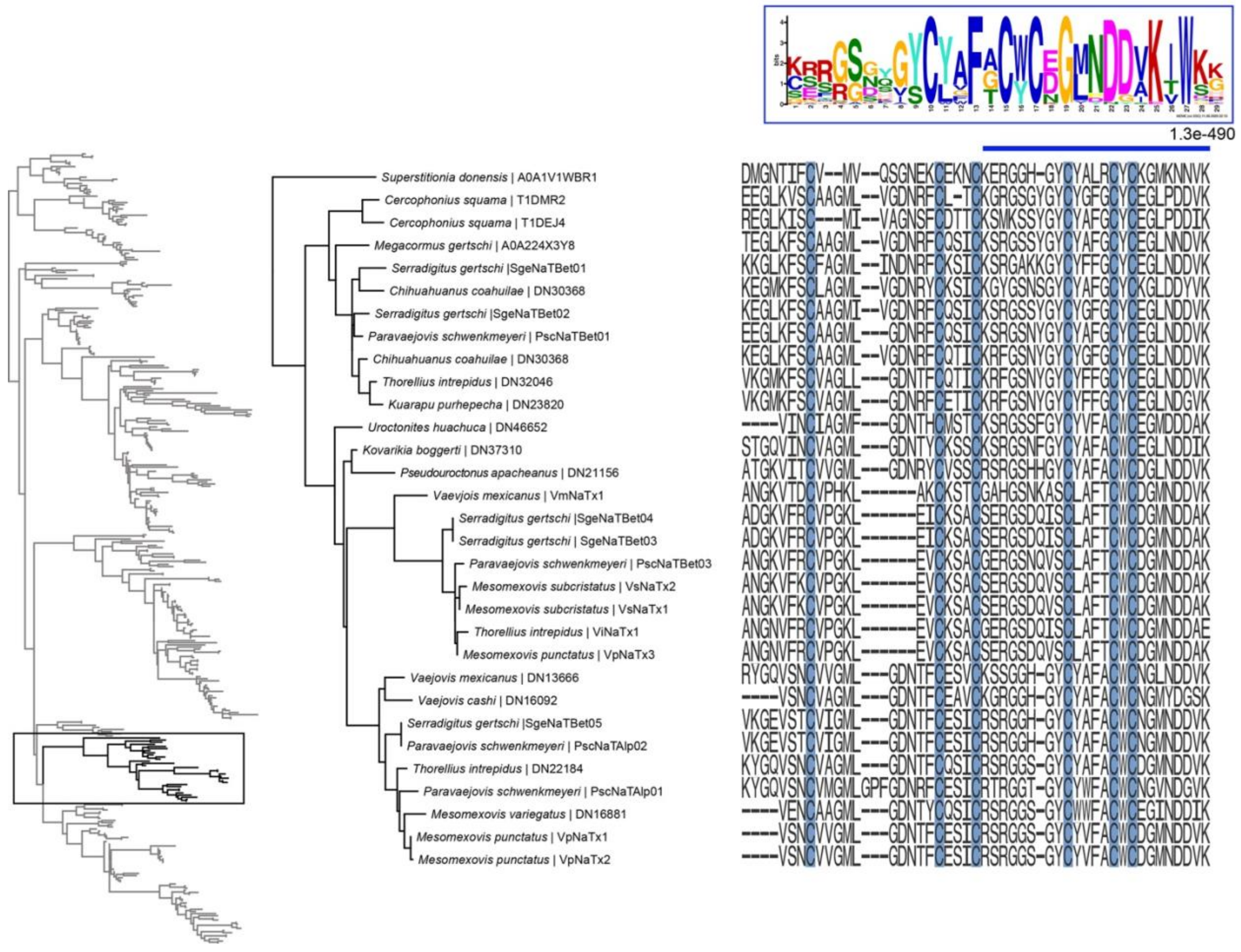

**Supplementary figure S13.** Left: Pruned gene tree of the Cn2-like clade unique to iurids. Right: Multiple sequence alignment of the mature peptide showing the repetitive motifs found using Multiple Em for Motif Elicitation (MEME).

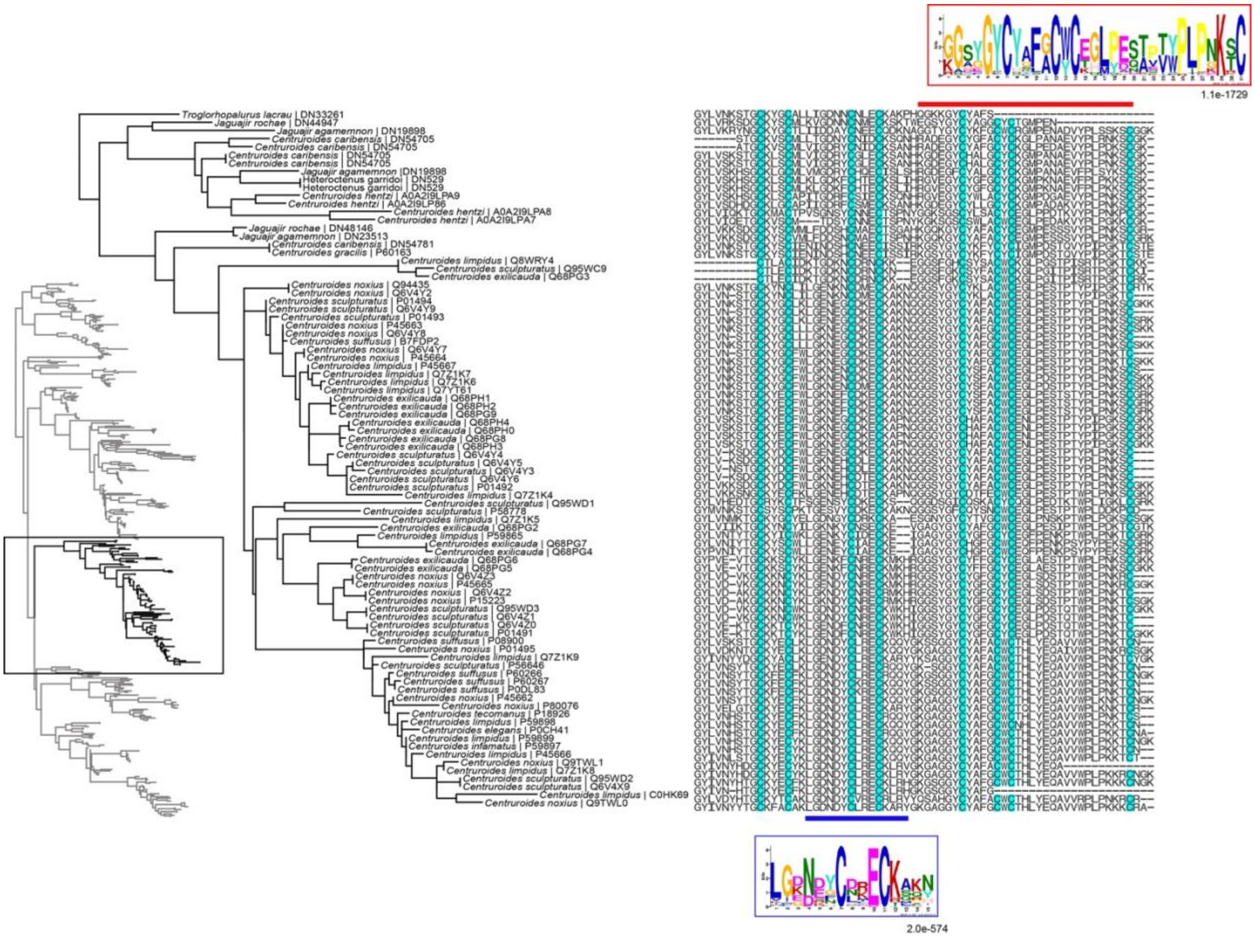


**Supplementary figure S14.** Left: Pruned gene tree of the Cn2-like clade unique to the “*Tityus*” group, with emphasis on the genus *Centruroides*. Right: Multiple sequence alignment of the mature peptide showing the repetitive motifs found using Multiple Em for Motif Elicitation (MEME).

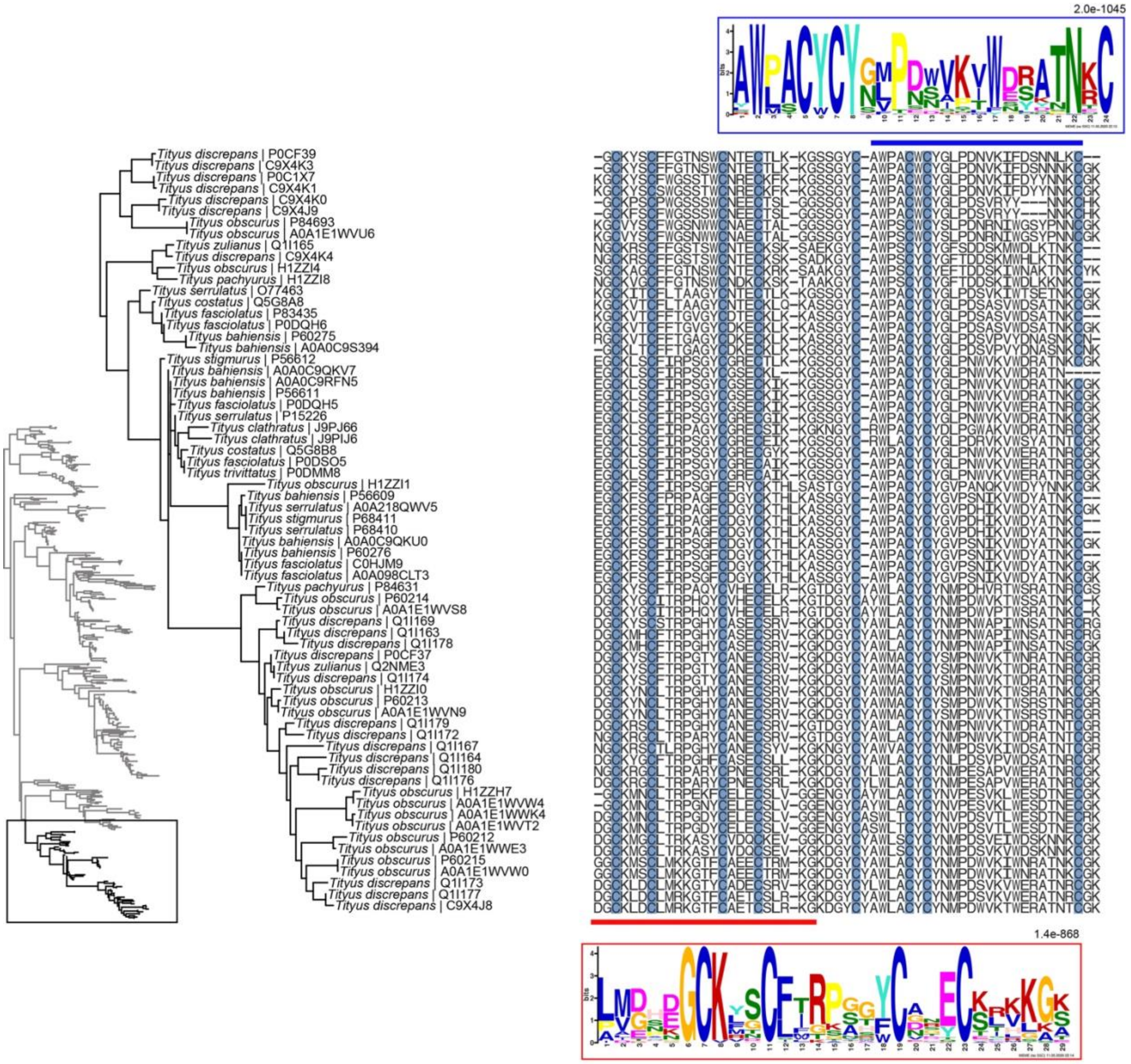


**Supplementary figure S15.** Left: Pruned gene tree of the Cn2-like clade unique to the genus *Tityus*. Right: Multiple sequence alignment of the mature peptide showing the repetitive motifs found using Multiple Em for Motif Elicitation (MEME).

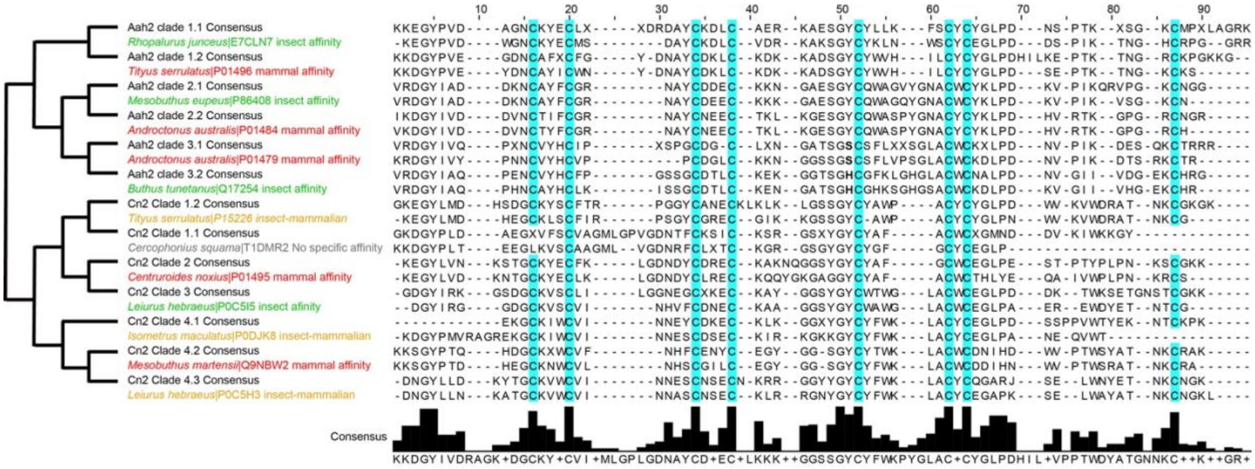


**Supplementary figure S16.** Left: Comparative analysis of the consensus clades containing at least one peptide sequence with known function shown as function of their phylogenetic relationships. Right: Multiple sequence alignment of the consensus sequence (in black) and a representative sequence from that clade with known function. Green text: insect affinity; orange text: insect and mammal affinity; red text: mammal affinity.

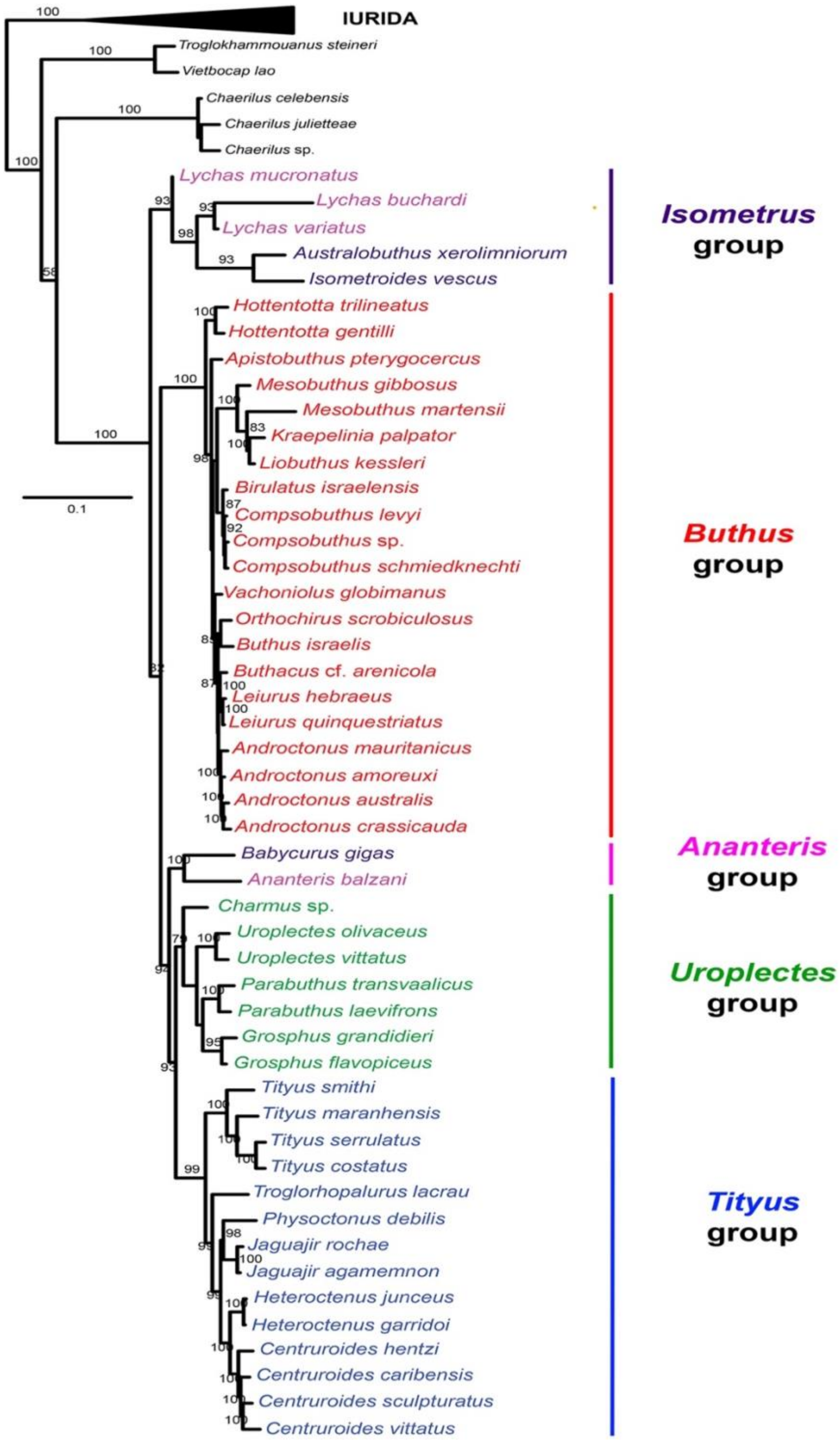


**Supplementary figure S17.** Maximum likelihood tree topology of Buthida recovered from the analysis of a combined dataset consisting of 424 genes (Matrix 2) and 2,692 nucleotides corresponding to three genes (18S rRNA, 16S rRNA, and COI). Color coding indicates species groups delimited by Fet et al. (2005).
